# Structural variation and DNA methylation shape the centromere-proximal meiotic crossover landscape in Arabidopsis

**DOI:** 10.1101/2023.06.12.544545

**Authors:** Joiselle B. Fernandes, Matthew Naish, Qichao Lian, Robin Burns, Andrew J. Tock, Fernando A. Rabanal, Piotr Wlodzimierz, Anette Habring, Robert E. Nicholas, Detlef Weigel, Raphael Mercier, Ian R. Henderson

**Affiliations:** Department of Plant Sciences, University of Cambridge, Cambridge, CB2 3EA, United Kingdom; Department of Chromosome Biology, Max Planck Institute for Plant Breeding Research, D-50829 Cologne, Germany; Department of Molecular Biology, Max Planck Institute for Biology, Tübingen, D-72076 Tübingen, Germany

**Keywords:** Centromeres, meiosis, crossover, recombination, DNA methylation, Arabidopsis.

## Abstract

**Background:** Centromeres load kinetochore complexes onto chromosomes, which mediate spindle attachment and allow segregation during cell division. Although centromeres perform a conserved cellular function, their underlying DNA sequences are highly divergent within and between species. Despite variability in DNA sequence, centromeres are also universally suppressed for meiotic crossover recombination, across eukaryotes. However, the genetic and epigenetic factors responsible for suppression of centromeric crossovers remain to be completely defined.

**Results:** To explore the centromere-proximal recombination landscape, we mapped 14,397 crossovers against fully assembled *Arabidopsis thaliana* genomes. *A. thaliana* centromeres comprise megabase-scale satellite repeat arrays that load nucleosomes containing the CENH3 histone variant. Each chromosome possesses a structurally polymorphic 3-4 megabase region where crossovers were absent, that includes the satellite arrays, flanked by 1-2 megabase low-recombination zones. The recombination-suppressed regions are enriched for Gypsy/Ty3 retrotransposons, and additionally contain expressed genes with high genetic diversity that initiate meiotic recombination, yet do not crossover. We mapped crossovers at high-resolution in proximity to *CEN3*, which resolved punctate centromere-proximal hotspots that overlapped gene islands embedded in heterochromatin. Centromeres are densely DNA methylated and the recombination landscape was remodelled in DNA methylation mutants. We observed that the centromeric low-recombining zones decreased and increased crossovers in CG (*met1*) and non-CG (*cmt3*) mutants, respectively, whereas the core non recombining zones remained suppressed.

**Conclusion:** Our work relates the genetic and epigenetic organisation of the *A. thaliana* centromeres and flanking pericentromeric heterochromatin to the zones of crossover suppression that surround the CENH3-occupied satellite repeat arrays.

## Background

Meiosis is a specialized eukaryotic cell division where a single round of DNA replication is coupled to two rounds of chromosome segregation, to produce haploid gametes [1,2]. During the first meiotic division, homologous chromosomes physically pair and undergo recombination that can result in reciprocal genetic exchange, termed crossover [1–3]. Meiotic recombination and independent chromosome segregation increase genetic diversity by reshuffling parental genomes into the gametes [1,2]. Meiotic recombination is initiated via DNA double strand breaks (DSBs) catalyzed by SPO11 complexes [2,4,5]. SPO11-dependent DSBs are resected to form single-stranded DNA, which can then mediate strand invasion of a sister or homologous chromosome and be repaired as a reciprocal crossover, or non reciprocal gene conversion [2,4,5]. Two major pathways, termed Class I and Class II, are required for crossover formation in plants [2]. Class I crossovers are the numerical majority and also show the phenomenon of interference, where double crossovers are more widely spaced than expected at random, whereas Class II events do not show interference [1,2]. Meiotic recombination occurs during prophase-I, when replicated homologs are physically associated via a chromosome axis and the synaptonemal complex, which provide the physical context for recombination [1–3].

Meiotic recombination frequency is highly variable within eukaryotic genomes, and kilobase scale hotspots of both DSBs and crossovers exist in plants, animals and fungi, whose locations are defined by a combination of DNA sequence and epigenetic information [6,7]. Conversely, other genomic regions are strongly crossover-suppressed, including the centromeres, repetitive heterochromatin, mating-type loci and sex chromosomes [8–11]. It has been proposed that suppression of crossovers within and around centromeres is beneficial, as proximal exchanges are associated with aneuploidy in fungi and animals, including trisomy in humans [12–15]. In addition, crossovers may lead to non-allelic exchanges in repeat regions, with the potential to cause deleterious structural change [11].

The centromeres function to assemble the kinetochore complex, which mediates chromosome attachment to spindle microtubules, during mitotic and meiotic cell divisions [16]. Centromere DNA sequences are loaded with nucleosomes containing the CENH3/CENP-A histone variant, which assemble the kinetochore [17,18]. Despite a conserved role in CENH3/CENP A loading, centromere DNA sequences are highly divergent within and between species [19–21], ranging from a ∼120 base pair sequence in budding yeast, to megabase-scale satellite repeat arrays in plants and vertebrates [8,21–23]. Although eukaryotic centromeres are composed of diverse DNA sequences, all centromeres show meiotic crossover suppression that spreads into flanking regions, over distances of kilobases to megabases [8,10,12,22]. However, the genetic and epigenetic features that regulate centromere-proximal recombination are incompletely understood.

Long-read DNA sequencing technologies, including PacBio HiFi and Oxford Nanopore, have allowed complete assembly of complex repeat regions [22–24]. For example, long-read DNA sequencing led to the assembly of the *Arabidopsis thaliana* centromeres, which comprise megabase-scale arrays of a 178 bp tandem repeat (*CEN178*) that are the site of CENH3 loading [21,22,24]. Plant and animal centromeres are often densely cytosine methylated, although the specific pattern varies between species [22,23,25]. For example, the CENP-A occupied regions of human α-satellite centromere arrays show CG-context DNA hypomethylation [26]. In contrast, the Arabidopsis CENH3-enriched regions are densely CG methylated, but hypomethylated in the CHG context [22]. In Arabidopsis, CG and non-CG context DNA methylation are maintained by distinct methyltransferase enzymes; MET1 for CG, and CMT2, CMT3 and DRM2 for non-CG [27]. The Arabidopsis pericentromeric regions are dominated by transposable elements and are also enriched for heterochromatic chromatin marks including H3K9me2, H3K27me1, H2A.W6, H2A.W7, and the meiotic cohesin REC8 [28–30]. Using complete maps of the Arabidopsis genome, we sought to investigate how genetic and epigenetic information shape the crossover landscape in proximity to the centromeres.

We mapped 14,397 crossovers genome-wide, against complete assemblies of the Arabidopsis Col and Ler accessions, and precisely identified zones of centromere-proximal suppressed recombination. The crossover-suppressed zones contain structurally variable satellite repeat arrays that are densely DNA methylated and load CENH3 nucleosomes, which we propose exert a joint effect on the recombination landscape. Low-recombining zones flank the centromeres and contain expressed genes that show elevated genetic diversity, with a range of housekeeping and environment-response functions. These centromere-proximal genes show evidence for meiotic recombination initiation, but not crossovers, indicating that repair steps downstream of DSB formation are inhibited. Using a fluorescence-based selection strategy, we fine-mapped 913 crossovers in proximity to *CEN3* and observed punctate recombination hotspots that overlap gene islands embedded in pericentromeric heterochromatin. We additionally mapped 962 and 1,033 *CEN3*-proximal crossovers in mutants defective in maintenance of CG (*met1*), or CHG sequence context (*cmt3*) DNA methylation. Centromere-proximal crossovers decreased and increased in *met1* and *cmt3*, respectively, and fine-scale remodelling of the recombination landscape was observed, although the satellite arrays remained crossover-suppressed. Our maps provide functional insight into the genetic and epigenetic factors that shape recombination in proximity to the Arabidopsis centromeres.

## Results

### The landscape of centromere-proximal crossover frequency in Arabidopsis

The Arabidopsis Col-0 (hereafter Col) and Ler-0 (hereafter Ler) accessions are of Eurasian origin and show ∼0.5% difference between shared DNA sequences in the chromosome arms [31]. Short-read sequencing of F2 or backcross progeny from Col/Ler F1 parents has been used to map meiotic crossovers [9,32–36]. As the Col and Ler centromere sequences have been fully assembled using long-read DNA sequencing [21,22], we sought to utilise these genome maps with the available crossover data to examine centromere-proximal recombination. For analysis, we combined 1,009 crossovers mapped from a Col/Ler female BC1 population, 978 crossovers from a male BC1 population and 12,410 from an F2 population, giving 14,397 crossovers in total and representing 3,613 meioses [9,33]. For crossover mapping, we used a refined set of single nucleotide polymorphisms (SNPs) that had been filtered for quality, and Mendelian segregation ratios in recombinant populations (**Fig. S1A**). Against the Col-CEN assembly, the average filtered Col/Ler SNP density was 2.56 per kb, and crossovers were resolved to a median width of 3,992 bp (**Fig. S1A**). Following short-read alignment from each individual library, the refined Col/Ler SNPs were genotyped and a sliding window approach used to map crossovers (**Fig. S1A**) [33–35].

To define zones of centromere-proximal recombination suppression, we tallied crossovers in 10 kb windows and defined: (i) the Non-Recombining Zones (NRZs) as contiguous centromeric regions with an absence of crossovers, and (ii) the Low-Recombining Zones (LRZs) as the flanking windows within 1 cM of each NRZ boundary (**Fig. 1, Fig. S1C** and **Table S1**). 1 cM windows were selected to define the LRZs based on genetic map length, rather than physical distance, on the different chromosomes. In total, the NRZs span 17.8 Mb, and are flanked by 10.8 Mb of LRZs (**Fig. 1, Fig. S1C** and **Table S1**). The mean density of SNPs in the LRZs (3.85 SNPs/kb) was higher than in the chromosome arms (2.80 SNPs/kb), whereas it was lower in the NRZs (0.75 SNPs/kb), meaning we are not limited by SNPs for detection of LRZ crossovers. The lower SNP density in the NRZs will reduce the precision of mapping individual crossovers, but not the ability to detect them. The LRZs comprise 10 cM in total with a recombination rate of 0.93 cM/Mb, compared to 3.80 cM/Mb in the chromosome arms. The majority of the NRZs are composed of the *CEN178* satellite arrays (13.2 Mb or 74%), and additionally contain mitochondrial genome insertions, *5S* rDNA, telomere and *CEN159* repeat arrays (**Fig. 1**, **Fig. S1** and **Table S1**). Within each *CEN178* satellite array, 1.4-1.6 Mb regions show CENH3 log2(ChIP/Input) enrichment scores >2, indicating kinetochore locations (**Fig. 1** and **Table S1**). We conclude that centromeric crossover inhibition spreads significantly beyond the boundaries of the CENH3-occupied regions (mean=1.76 Mb), and *CEN178* satellite arrays (mean=2.54 Mb), with the joint LRZ-NRZ spanning on average 5.72 Mb per chromosome (**Fig. 1** and **Table S1**).

**Figure 1.**
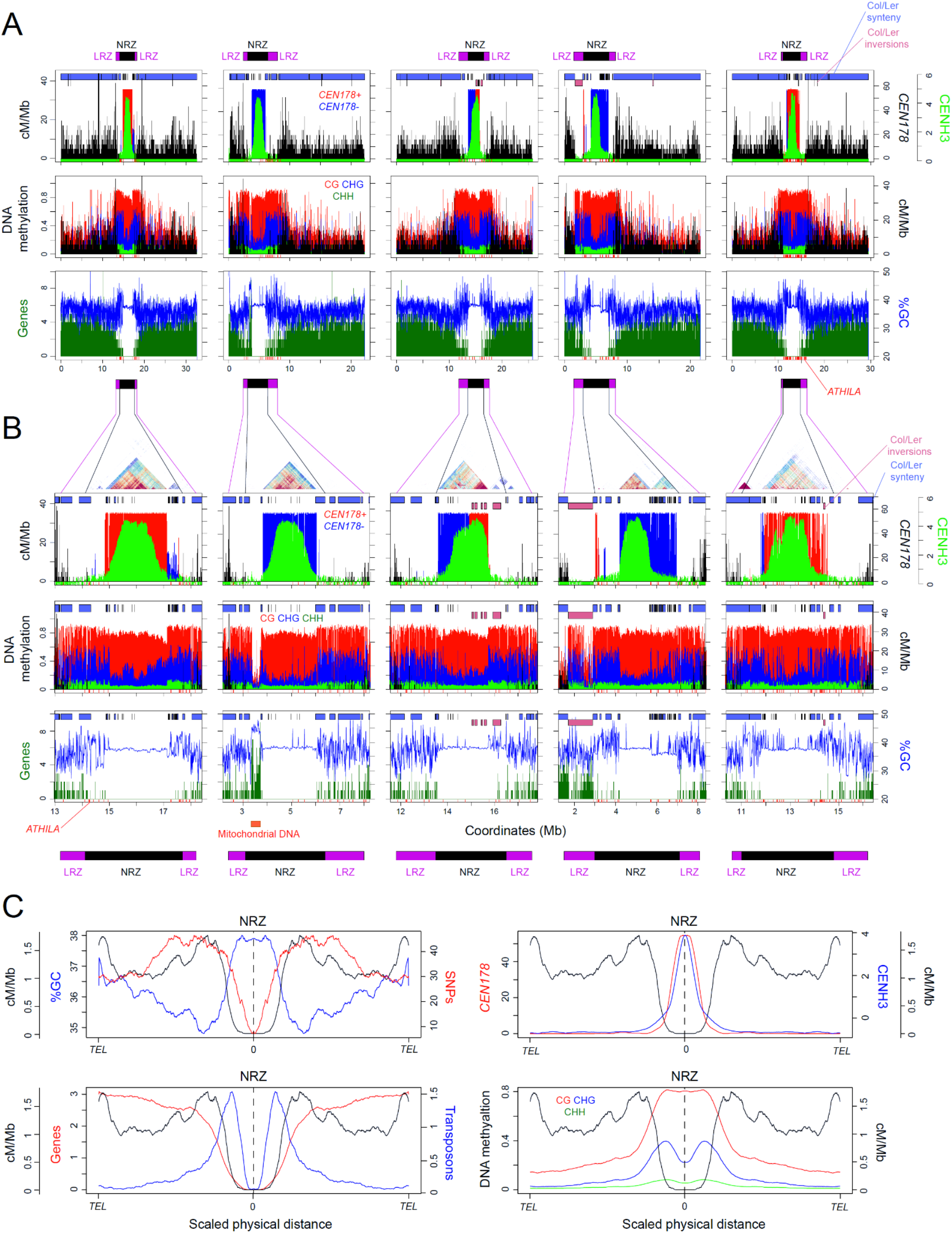
Zones of centromere-proximal crossover suppression in Arabidopsis. **A.** In the first row, Col/Ler crossover frequency (cM/Mb) is plotted against the Col-CEN assembly in 10 kb windows (black). *CEN178* satellite density (red=forward & blue=reverse strand) and CENH3 ChIP-seq enrichment (green, log_2_[ChIP/input]) are plotted in the same windows. Locations of the non-recombining zones (NRZ, black), and low-recombining zones (LRZ, purple), are indicated above. Shaded blocks at the top of the plots indicate regions of Col/Ler synteny (blue) and inversions (pink) mapped by SYRI [40]. In the next row, the proportion of DNA methylation mapped from ONT reads is plotted in 10 kb windows for CG (red), CHG (blue) and CHH (green) sequence contexts. Beneath, the density of genes (green) is plotted alongside %GC content (blue) per 10 kb. *ATHILA* retrotransposon insertions are indicated as red x-axis ticks. **B.** As for A, but showing a zoom of the NRZ and LRZ regions. Plot annotations are the same, apart from a StainedGlass sequence identity heatmap is positioned over the plots [79], and NRZ-LRZ positions are shown beneath. **C.** Quantification of genomic features plotted along chromosome arms that were proportionally scaled between telomeres (*TEL*) and NRZ midpoints. Data analysed were gene, transposon and *CEN178* density per 10 kb, CENH3 log_2_(ChIP/input), %GC base composition, DNA methylation, and crossovers (cM/Mb).

The LRZs and NRZs are densely DNA methylated in CG, CHG and CHH sequence contexts, although the CENH3-enriched regions within the NRZs show relative depletion of CHG context methylation (**Fig. 1**). CHG depletion in the centromeres is proposed to be a consequence of CENH3 being unable to sustain H3K9me2 histone methylation, which is required to maintain CHG context DNA methylation in Arabidopsis [22,27]. The LRZs are strongly enriched for heterochromatic histone modifications H3K9me2, H3K27me1, H2A.W6 and H2A.W7, although similar to CHG DNA methylation, these marks are relatively depleted within NRZ-CENH3 regions (**Fig. S2**) [28,37]. Interestingly, H2A.W7 showed a stronger depletion in the CENH3-enriched regions compared to H2A.W6 (**Fig. S2**). Both LRZs and NRZs have significantly higher GC base content (37.6% and 38.2%), compared to the chromosome arms (35.8%) (Wilcox test *P*=0.0079) (**Fig. 1**). ChIP-seq enrichment of REC8-cohesin, and the HORMA domain protein ASY1, which are components of the meiotic chromosome axes, are strongly enriched in the LRZs and NRZs, and to a lesser extent within the NRZ-CENH3 regions (**Fig. S3**) [28,38]. We detected peaks of SPO11-1-oligos, a marker of meiotic DNA double strand breaks, within the LRZs, which correlated with crossovers in a subset of cases (**Fig. S3**) [39]. SPO11-1-oligo peaks observed in the absence of crossovers may reflect initiation of meiotic DSBs, but repression of downstream crossover repair by centromeric chromatin states, or structural polymorphism.

To explore patterns of structural polymorphism, we used Synteny and Rearrangement Identifier (SyRI) to identify syntenic regions between the Col and Ler genomes, in addition to rearrangements (**Fig. 1A-1B**) [40]. The LRZs-NRZs show disrupted synteny between Col and Ler, consistent with structural polymorphism contributing to centromere-proximal crossover suppression (**Fig. 1A-1B**). This includes inversion of the chromosome 3 *CEN178* array and flanking sequences, and a large pericentromeric ‘knob’ inversion adjacent to *CEN4* (**Fig. 1A-1B**) [22,41]. We compared the structure of the satellite arrays between Col and Ler, for each chromosome, and observed significant structural polymorphism, despite them being composed of the same *CEN178* repeats (**Fig. 1** and **Fig. S4**). For example, both *CEN1* and *CEN2* are larger (2.67 & 2.91 vs 1.91 & 1.51 Mb) and more repetitive in Col (**Fig. S4**). Col and Ler *CEN3* show array inversions, but the Ler centromere is larger and consists of discontinuous islands of repeats on its left boundary (**Fig. S4**). In both accessions, *CEN4* consists of two adjacent, yet distinct, *CEN178* arrays, with the left array being CENH3 occupied and more divergent in sequence between the accessions (**Fig. S4**). In contrast*, CEN5* is larger and more repetitive in Ler (3.37 vs 2.66 Mb) (**Fig. S4**). The extensive *CEN178* repeat array polymorphisms between the Col and Ler genomes could directly contribute to NRZ crossover suppression, in addition to the effects of epigenetic information.

### Gene and transposon content of the centromeric recombination-suppressed zones

The LRZs and NRZs contain 542 and 132 genes, respectively, excluding the mitochondrial genes (n=106) located adjacent to *CEN2* [42]. The LRZs and NRZs have lower gene density (50 and 7 genes/Mb, respectively), compared to the chromosome arms (268 genes/Mb). Despite the NRZ and LRZ being heterochromatic, 42% and 48% of the contained genes showed evidence of expression from RNA-seq data, respectively (**Fig. 2A** and **Table S2**) [43]. NRZ and LRZ genes included those with house-keeping and environment-response annotation, genes with genetically defined roles (e.g. *ARP6*, *CLASSY1*, *MBD12* and *OSCA1*), and with putative roles in immunity (e.g. defensins, *TIR-NBS14*, and WRKY transcription factors) (**Table S2**). We also observed eight gene clusters, comprised of at least four orthologs, that encode calcium-dependent kinases, ankyrin-repeats, nucleotide transporters, receptor-kinases, ubiquitin, F-box proteins, sulfoxide reductases and lipid-transfer proteins (**Table S2**).

**Figure 2.**
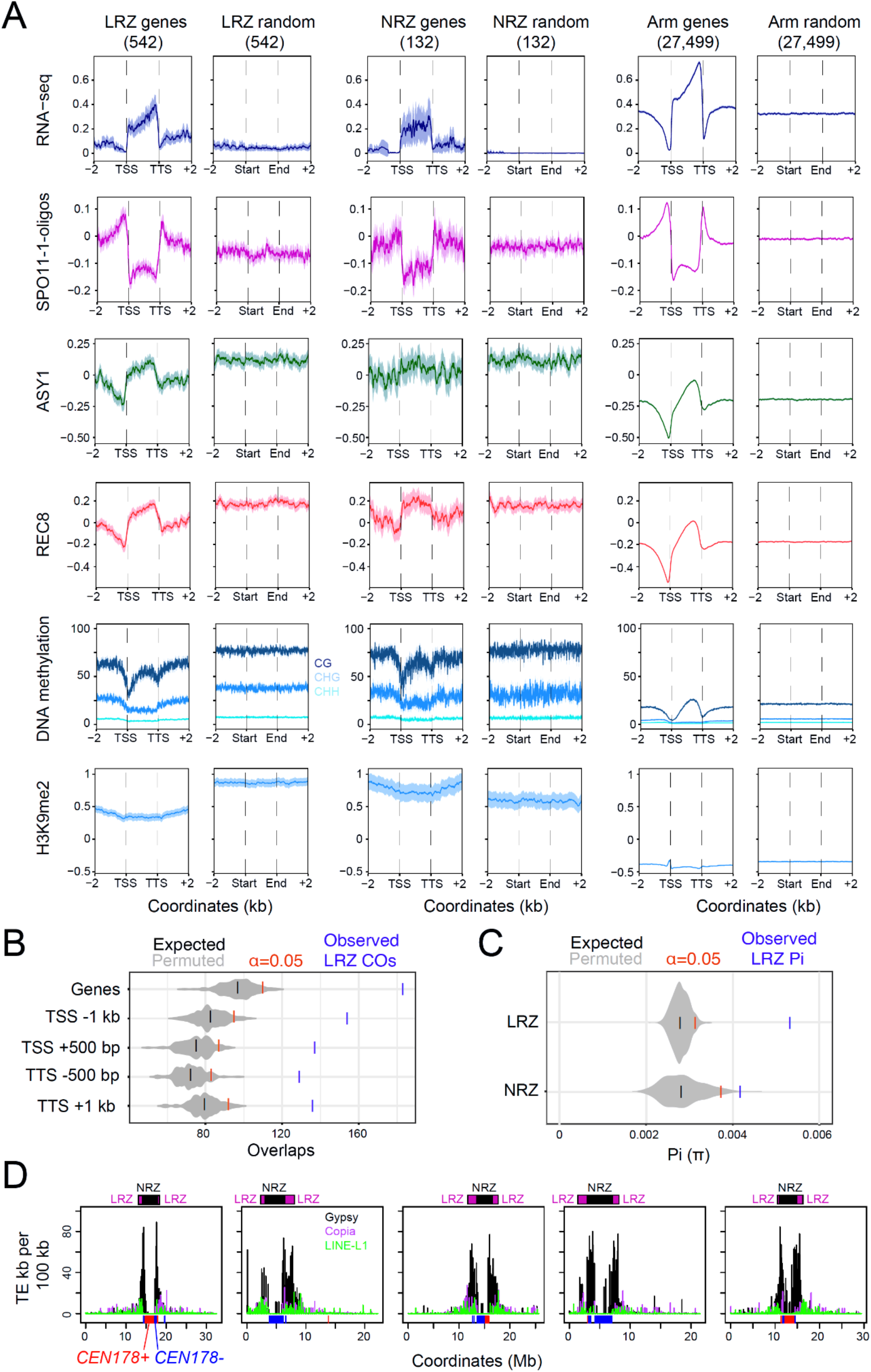
Gene and transposon content of the Arabidopsis LRZs and NRZs. **A.** Metaprofiles of leaf RNA-seq (blue, Log_2_[RNA-seq (TPM)] [86], SPO11-1-oligos (pink, Log_2_[SPO11-1-oligos/genomic DNA]) [39], ASY1 (green, Log_2_[ChIP-seq/input]) [38], REC8 (red, Log_2_[ChIP-seq/input]) [28], DNA methylation (%, CG=dark blue, CHG=blue, CHH=light blue) [22], and H3K9me2 (blue, Log_2_[ChIP-seq/input]) [28], across genes located in the chromosome arms (n=27,499), the LRZs (n=542) and the NRZs (n=132) and in 2 kb flanking regions. For each gene set, the same number of random windows of the same widths were compared within the same regions. Plot ribbons denote 95% confidence intervals for windowed values. **B.** Observed number of LRZ crossovers overlapping the listed gene features are shown (blue), compared to 10,000 sets of randomly positioned loci of the same number and width distribution as the LRZ crossovers. The α=0.05 significance thresholds are indicated (red), and the means of the permuted sets of loci (black) (*P*-values from all comparisons were <0.0001). **C.** The observed median Pi (π) value for genes located in the LRZs (n=336) and NRZs (n=58) (blue), compared to 1,000 sets of randomly chosen genes in the chromosome arms (grey). α=0.05 significance thresholds are indicated (red), and the medians of the permuted loci sets (black). *P*-values for both comparisons were <0.0001. Pi was calculated using 1,001 Genomes Project SNPs [46]. **D.** Plots of retrotransposon density per 100 kb along the Col-CEN assembly, showing Gypsy (black), Copia (purple) and LINE (green) superfamilies. The locations of the LRZs (purple) and NRZs (black) are indicated above the plots, and the *CEN178* satellite arrays (red, blue) are indicated along the x-axis.

The LRZ and NRZ genes showed comparable accumulation of SPO11-1-oligos in their promoters and terminators, compared to chromosome arm genes (**Fig. 2A**) [39]. Chromosome arm gene bodies are enriched for REC8-cohesin and ASY1, which was also observed for LRZ and NRZ genes, although levels were higher in the LRZ/NRZ genes (**Fig. 2A**) [28,38]. DNA methylation levels were higher in the NRZ and LRZ genes compared to the chromosome arms, yet they retained a typical profile of relative DNA hypomethylation in promoters and terminators and CG context methylation within their open reading frames (**Fig. 2A**). One striking difference between the NRZ/LRZ genes and those in the chromosome arms, is greater enrichment of the heterochromatic H3K9me2 histone modification (**Fig. 2A**) [28]. As H3K9me2 has been associated with crossover suppression in Arabidopsis [44,45], we propose this contributes to inhibition of crossover repair, despite significant SPO11-1-oligos forming around NRZ and LRZ genes (**Fig. 2A**).

Although LRZ crossovers are relatively suppressed compared to the chromosome arms, we investigated whether they locally overlapped genes. We compared observed overlaps between crossovers and LRZ genes, with overlaps of randomly positioned loci. For statistical comparison, 10,000 permuted randomly positioned sets within the LRZa of the same number and widths as the crossovers were used (**Fig. 2B**). We observed that LRZ crossovers significantly overlapped with genes and their upstream and downstream regions (all tests *P*=9.99⨉10^-5^) (**Fig. 2B**). We also analysed transposon content within the NRZs and LRZs, and observed strong enrichment of Gypsy/Ty3 LTR class retrotransposons (151.0 and 73.2 per Mb, respectively), compared to the chromosome arms (12.0 per Mb) (**Fig. 2D**). This implies that Gypsy/Ty3 retrotransposons may actively integrate into the LRZs and NRZs, or their insertions have been selected against in the chromosome arms. We repeated permutation tests for crossover overlap with LRZ transposons and observed significant overlap with Helitron transposons, and significant non-overlap with Gypsy/Ty3 retrotransposons (both *P*=<9.00⨉10^-5^), consistent with previous positive and negative associations of these families with meiotic DSBs, respectively [39].

We examined NRZ and LRZ gene diversity compared to the chromosome arms. We calculated pairwise diversity (Pi, π) for genes, using SNPs from 1,135 Arabidopsis accessions [46]. The SNPs were masked for repeat sequences and we required that at least half of the gene had sequencing coverage across the 1,135 accessions to be included. We also excluded genes that overlapped Col/Ler inversions, and the mitochondrial genome insertions on chromosome 2. After filtering, we retained 336 LRZ and 58 NRZ genes for analysis, calculated median Pi, and compared to 1,000 permutations of the same numbers of genes from the chromosome arms (**Fig. 2C**). The observed median value of Pi for genes located in the LRZs and NRZs, were significantly higher than the permuted sets from the chromosome arms (both *P*=<0.0001) (**Fig. 2C**). Further work will be required to ascertain the relative contributions of recombination, demography, mutation rate and selection to elevated LRZ and NRZ gene diversity.

### Fine-mapping of meiotic crossovers in proximity to *CENTROMERE3*

As centromere-proximal crossovers are rare, we designed a fluorescence-based selection strategy to enrich for recombination events and perform fine-mapping (**Fig. 3A**). In Arabidopsis, linked hemizygous T-DNAs (fluorescence-tagged lines, FTLs) expressing different colours of fluorescent protein in the pollen or seed can be used to quantify and map intervening crossovers (**Fig. 3A**) [47,48]. We selected the *CTL3.9* FTL, generated in the Col accession, to investigate crossover frequency in proximity to *CEN3* (**Fig. 3**) [48]. Against the Col-CEN assembly, the *CTL3.9* T-DNAs define a 8.5 Mb interval, which includes the 2.14 Mb *CEN178* satellite repeat arrays (**Fig. 3**) [22]. As noted, *CEN3* in Col and Ler contain adjacent *CEN178* arrays on opposite strands (**Fig. 1** and **3**) [22]. The remainder of the *CTL3.9* interval is heterochromatic, containing numerous DNA and RNA transposable elements, yet shows increasing gene density towards the *CTL3.9* T-DNAs (**Fig. 3**).

**Figure 3.**
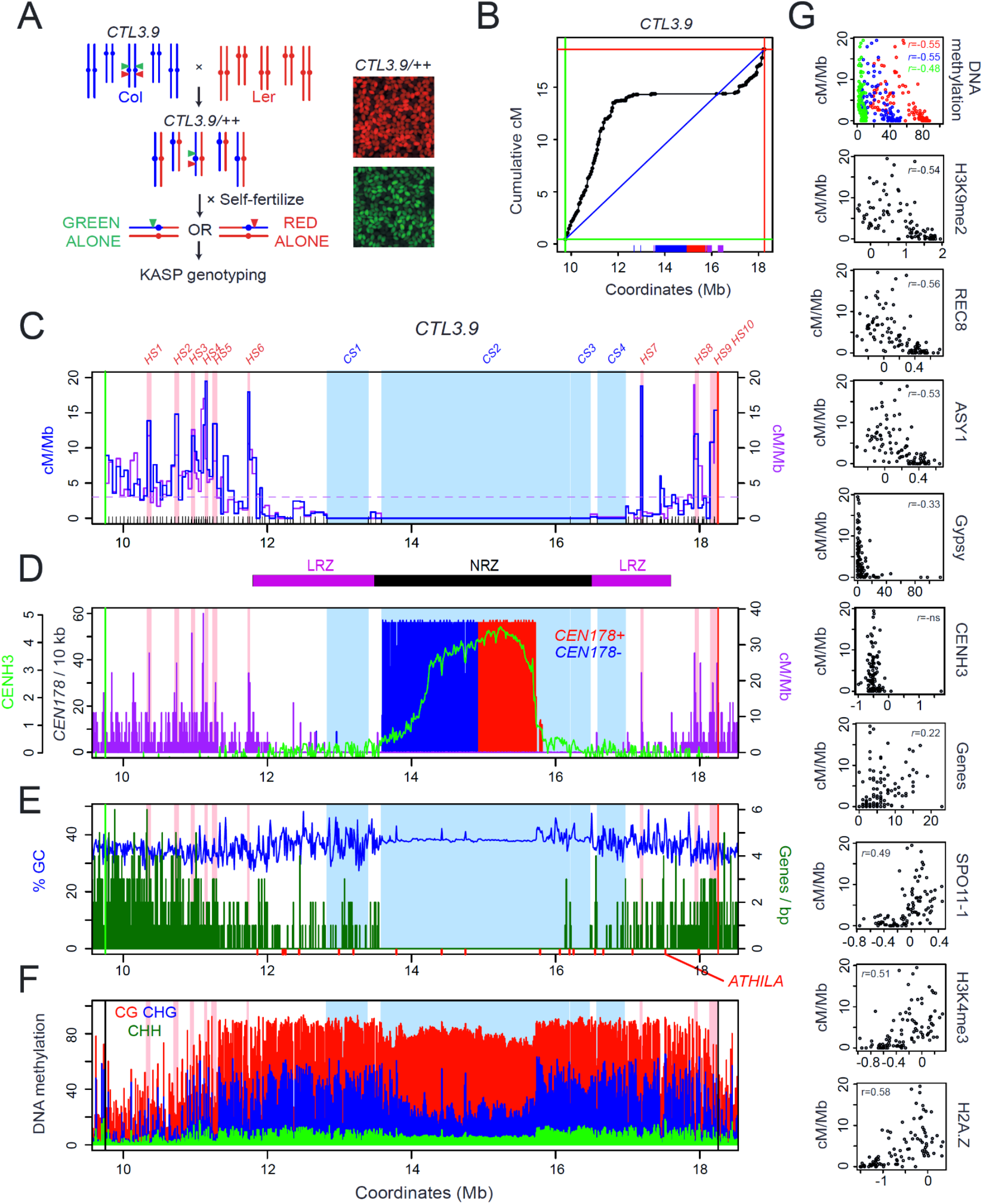
The fine-scale recombination landscape around Arabidopsis centromere 3. **A.** Genetic strategy to recover crossovers within the *CTL3.9* FTL interval. FTL T-DNAs encoding red and green fluorescent proteins are indicated by triangles. The parental chromosomes are from the Col (blue) and Ler (red) accessions. Fluorescent micrographs of *CTL3.9/++* segregating seed are shown to the right. **B.** Cumulative genetic map (centiMorgans, cM) relative to *CTL3.9* genomic coordinates plotted against the Col-CEN assembly derived from KASP genotyping of selected recombinant seed. The blue diagonal line shows a linear relationship, with the red and green lines showing the *CTL3.9* T-DNAs. The position of *CEN178* satellite repeats are shown as red (forward) and blue (reverse) ticks on the x-axis, in addition to *5S* rDNA (purple). **C.** Crossover frequency (centiMorgan per megabase, cM/Mb, blue) plotted within *CTL3.9* derived from KASP genotyping, and compared to measurements from mapping-by-sequencing (purple) for the same intervals. FTL T-DNAs are indicated by red and green vertical lines. Col/Ler KASP marker positions are indicated as x-axis ticks. The horizontal dotted line shows the genome average cM/Mb. Map intervals with significantly higher (*HOTSPOT, HS*) or lower (*COLDSPOT*, *CS*) crossover counts are shaded pink and blue, respectively. The black and purple bars beneath show the NRZ and LRZs. **D.** As for C, but the frequency of *CEN178* satellite repeats on forward (red) and reverse (blue) strandsper 10 kb is shown, overlaid with CENH3 log2(ChIP/input) enrichment (green) and GBS cM/Mb (purple). **E.** As for C, but showing gene density per 10 kb (green), overlaid with %GC content (blue). *ATHILA* retrotransposon positions are indicated by x axis ticks (red). **F.** As for C, but showing the proportion of DNA methylation in 10 kb windows for CG (red), CHG (blue) and CHH (green) contexts. **G.** cM/Mb values for *CTL3.9* map intervals are presented as scatter plots compared against DNA methylation, H3K9me2, REC8, ASY1 ChIP-seq, Gypsy/Ty3 transposons, CENH3 ChIP-seq, genes, SPO11-1-oligonucleotides, H3K4me3 and H2A.Z. Spearman’s correlation coefficient is printed inset, where significant.

Arabidopsis T-DNAs may be present as multi-copy insertions, or associated with chromosome rearrangements, in addition to being DNA methylated [49,50]. As both structural variation and DNA methylation suppress Arabidopsis crossovers [9,44,45], we sought to map the genetic and epigenetic state of the *CTL3.9* T-DNAs. We sequenced the *CTL3.9* line using Oxford Nanopore Technology (ONT) and performed genome assembly (**Fig. S5**). The *CTL3.9* genome assembly showed no evidence of large-scale structural rearrangements, compared to the parental Col line (**Fig. S5A**). The *CTL3.9* loci CG17 (green) and CR55 (red) comprise 12.7 and 5.1 kb insertions flanked by T-DNA border sequences (**Fig. S5C-S5D**). We used Deepsignal-plant to map patterns of DNA methylation over the *CTL3.9* T-DNAs, using our ONT data [51]. The T-DNAs were DNA methylated in CG, CHG and CHH sequence contexts, although at lower levels than flanking transposable elements (**Fig. S5E-S5F**). Although DNA methylation and structural hemizygosity of the FTL T-DNAs may cause local crossover suppression, these effects are likely to be limited relative to the size of the entire ∼8.5 Mb *CTL3.9* interval. Furthermore, inheritance of red and green fluorescence from *CTL3.9* hemizygotes conformed to the Mendelian expectation of ∼3:1 colour:non-colour, which is further consistent with the absence of rearrangements or significant transgene silencing (**Tables S3-S4**).

To map crossovers within *CTL3.9*, we crossed to the genetically polymorphic accession Ler. In wild type inbreds (Col/Col), *CTL3.9* has a genetic distance of 16.5 cM, equivalent to a recombination rate of 1.94 cM/Mb (**Table S3**), compared to the genome average of 3.03 cM/Mb. In Col/Ler F_1_ hybrids, *CTL3.9* crossover frequency significantly increased relative to Col/Col inbreds, with a genetic distance of 18.3 cM, equivalent to 2.15 cM/Mb (Wilcoxon test *P=*2.74⨉10^-6^) (**Table S4**). This is consistent with Arabidopsis hybrids showing higher crossover frequency than inbreds in other genetic intervals [52,53]. From *CTL3.9* hemizygous Col/Ler F_1_ plants, we selected progeny seed that showed either red or green fluorescence alone, consistent with a single crossover between the T-DNAs (**Fig. 3A**). We selected 444 red-alone and 464 green-alone seeds, which were sown and genotyped with an array of 94 Kompetitive Allele-Specific PCR (KASP) markers to genotype Col/Ler SNPs within *CTL3.9*, with an average inter-marker distance of 89.5 kb (**Fig. 3A** and **Table S10**). We mapped 913 crossover events in total within *CTL3.9* (**Table S5**). The genotypes of 908 plants were consistent with a single chromatid having one crossover and the other chromatid being non recombinant, while two plants each contained two single crossover chromatids (**Fig. S6**).

Crossovers were unevenly distributed within *CTL3.9*, and the recombination landscape was significantly correlated with crossovers mapped via whole-genome sequencing (*r*=0.719 *P*=<2.2⨉10^-16^) (**Fig. 3C** and **Table S5**). To define crossover hotspots and coldspots, we calculated the expected number of events per interval, assuming an even distribution, and compared this to observed values (**Table S6**). Observed and expected crossover counts for each interval were used in chi-square tests, followed by Bonferroni correction for multiple testing, to identify intervals that contained significantly higher or lower recombination. This approach identified four cold spot intervals (*CS1-CS4*), which occupy the central region of *CTL3.9*, and ten hotspots (*HS1-HS10*), which are distributed throughout the LRZs, with crossover rates in the range 10.8–19.5 cM/Mb (**Fig. 3C** and **Table S6**). Several hotspots, for example *HS6* and *HS7*, are located inside dense heterochromatin close to the NRZ boundaries (**Fig. 3C**). The *HS* intervals showed crossover rates comparable to previously mapped euchromatic hotspots in Arabidopsis [54–58]. We observed that compared with the coldspots, the hotspot intervals had significantly lower CG, CHG and CHH context DNA methylation, ASY1, CENH3, H3K9me2 and REC ChIP-seq enrichment, and significantly higher H3K4me3 and H2A.Z ChIP-seq enrichment, and SPO11-1-oligos (all Wicox tests <7.99⨉10^-3^) (**Fig. S7**). We further correlated chromatin states with cM/Mb in all *CTL3.9* map intervals and observed significant negative relationships with DNA methylation (CG, CHG and CHH contexts), H3K9me2, REC8 and ASY1 ChIP-seq, and positive relationships with SPO11-1-oligos, H3K4me3 and H2A.Z ChIP-seq (**Fig. 3G**). Overall, this is consistent with euchromatic marks promoting meiotic DSBs and crossovers, and heterochromatic marks repressing crossovers.

To explore how genetic and epigenetic polymorphism influence recombination, we projected our crossover data onto the Col and Ler genome assemblies (**Fig. 4**). The LRZs and NRZ showed high levels of structural polymorphism (**Fig. 4**), which may directly contribute to crossover suppression in the central cold spot intervals. We compared epigenetic maps of DNA methylation and CENH3 ChIP-seq enrichment generated for each Col and Ler genome assembly (**Fig. 4**) [21]. The Col and Ler centromeres both contain ∼1 Mb regions of CENH3 ChIP-enrichment within the centre of the main *CEN178* arrays, which are typified by reduced CHG context DNA methylation relative to the flanking pericentromeres (**Fig. 4**). This is further consistent with a combination of DNA sequence and chromatin factors determining centromere-proximal recombination in Arabidopsis.

**Figure 4.**
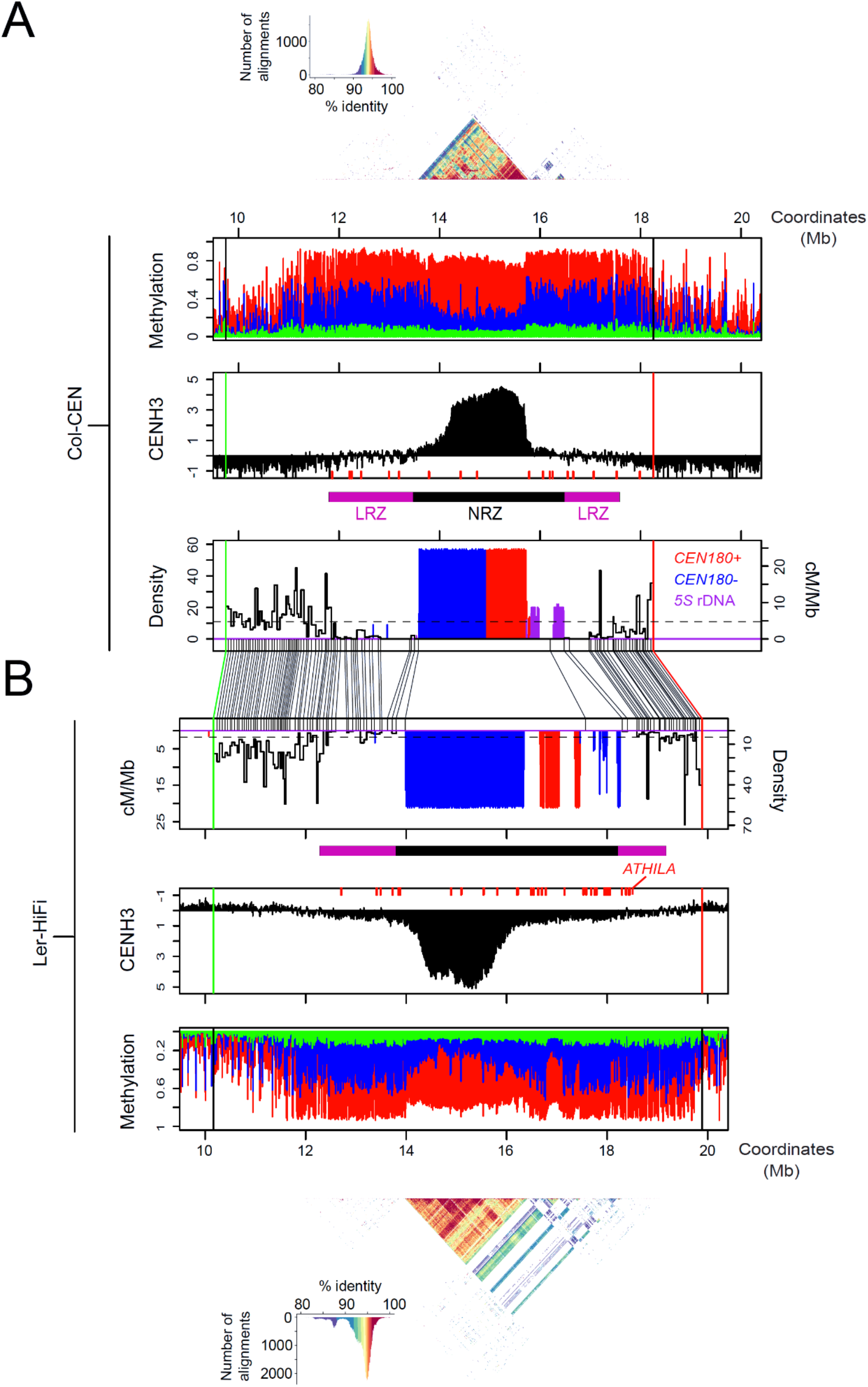
Genetic and epigenetic haplotypes of Col and Ler centromere 3 and the crossover recombination landscape. **A.** *CTL3.9* crossover frequency (cM/Mb) is plotted against the Col-CEN assembly, overlaid with a plot of the density of *CEN178* satellite repeats (red=forward, blue=reverse strand) and *5S* rDNA (purple). The positions of the *CTL3.9* fluorescent T-DNAs are shown by red and green vertical lines. Above this plot, the purple and black bars indicate the positions of the LRZs and NRZ. Above is a plot of CENH3 ChIP-seq enrichment (black), with *ATHILA* retrotransposons indicated by red x-axis ticks. Above this is a plot of the proportion of ONT-based DNA methylation in CG (red), CHG (blue) and CHH (green) sequence contexts. A StainedGlass sequence identity heat map is shown above the *CEN178* satellite arrays, together with a histogram indicating the colour scale associated with % identity values. **B.** As for A, but showing a mirrored version with data projected and aligned to the Ler-0 genome assembly. In the centre of the plot, the physical positions of KASP and T-DNA markers in the Col-CEN and Ler-HiFi assemblies are connected with lines between the x-axis.

### The centromere-proximal *HOTSPOT6* contains gene islands embedded in heterochromatin

*HOTSPOT6* (*HS6*) was centromere-proximal on the left arm and showed an elevated crossover rate (17.9 cM/Mb) (**Fig. 3C**). We sought to fine-map *HS6* crossovers (n=23) using additional markers (**Fig. 5A**). We designed four Col/Ler derived Cleaved Amplified Polymorphic Sequence (dCAPS) markers within *HS6* and genotyped each of the 23 samples that contained a crossover (**Fig. 5** and **Table S7**). Crossovers were observed throughout *HS6*, with highest rates towards the centromere-proximal boundary (**Fig. 5** and **Table S7**). Four genes are located within *HS6*, encoding a P-loop NTP hydrolase, COBRA-like2, an F-box protein, and a B12D transmembrane protein (**Fig. 5B**). Of these genes, *COBRA-like2* showed highest evidence of expression from RNA-seq data (**Fig. 5B**) [43]. The *HS6* genes are either methylated in CG context alone, or unmethylated, whereas the intervening regions contain multiple transposon sequences that are densely DNA methylated in all contexts (**Fig. 5B**). The gene regions were also distinguished by higher H3K27me3 and H2A.Z ChIP-seq enrichment (**Fig. 5B**) [37]. SPO11-1-oligos within *HS6* were denser in proximity to the genes and reduced within the heterochromatic repeats, whereas the opposite was true for REC8-cohesin and ASY1 ChIP-seq (**Fig. 5B**) [28,38]. This shows that *HS6* crossovers are associated with gene islands that are otherwise embedded in repetitive heterochromatin.

**Figure 5.**
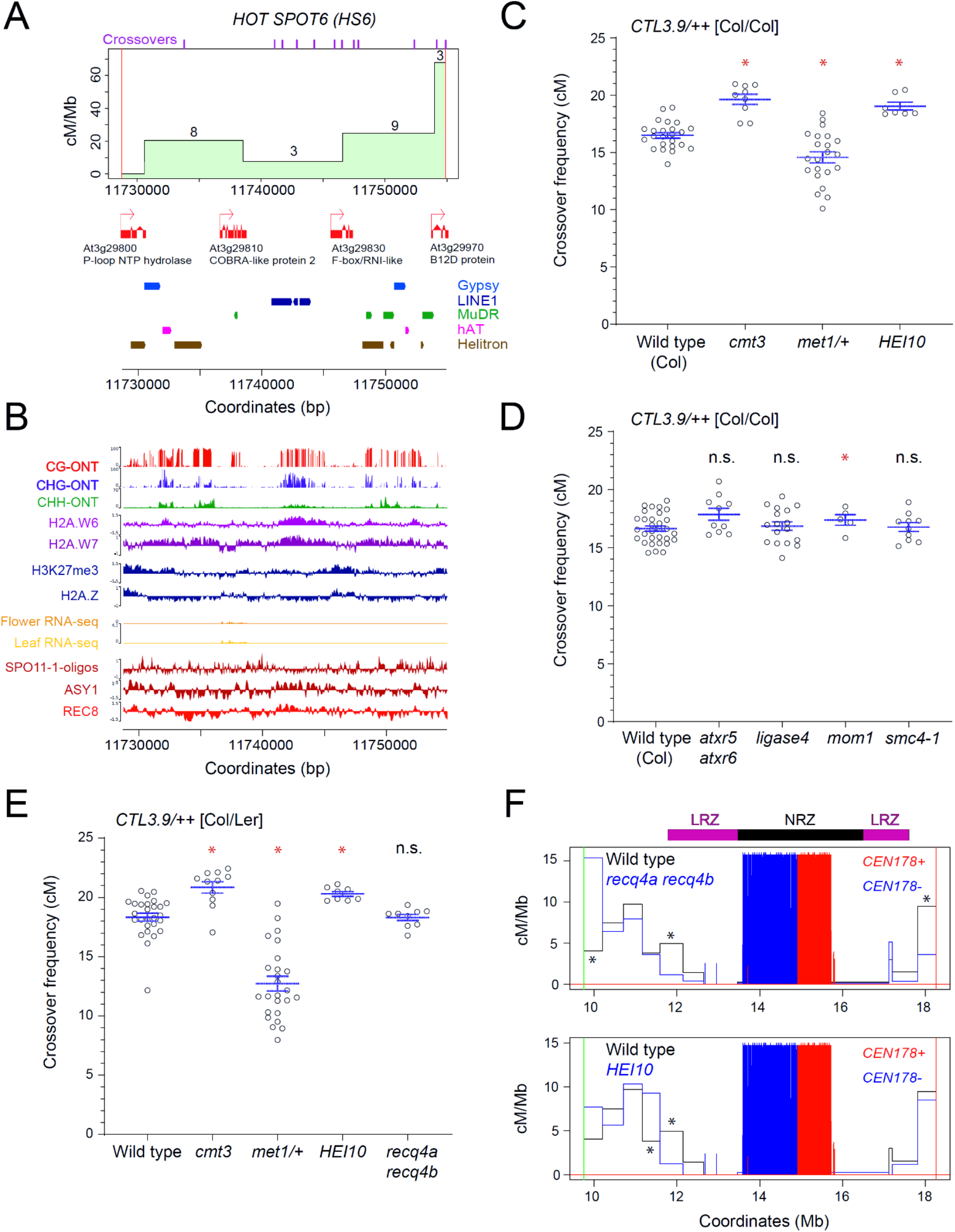
Genetic and epigenetic control of crossover frequency within *CTL3.9* and the *HS6* hot spot. **A.** Derived cleaved amplified polymorphic sequence (dCAPS) markers were used to map crossovers within *HOT SPOT6* (*HS6*) and calculate cM/Mb. The numbers printed above the plot line show the number of crossovers identified in each interval. Crossovers mapped by sequencing, as in Fig. 1, are indicated as purple ticks along the top axis. Gene annotation (red) is shown underneath, in addition to Gypsy/Ty3 (blue), LINE1 (dark blue), MuDR (green), hAT (pink) and Helitron (brown) transposon annotations. **B.** Plots of the *HS6* interval showing ONT-derived DNA methylation (%) in CG (red), CHG (blue) or CHH (green) sequence contexts, ChIP-seq enrichment (log_2_(ChIP/input)) for H2A.W6 (light purple), H2A.W7 (dark purple), H3K27me3 (blue), H2A.Z (blue), ASY1 (dark red) and REC8 (red), RNA-seq from floral (orange) and leaf (yellow) tissue, and SPO11-1-oligos (dark red). **C.** *CTL3.9* crossover frequency (cM) in wild type, *cmt3*, *met1-3/+* and *HEI10*, in an otherwise Col/Col homozygous background. Measurements from individuals are shown as circles. Horizontal blue lines represent the mean and the standard error of the mean. Red stars indicate samples which are significantly different from wild type using Wilcoxon tests, whereas ‘n.s.’ indicates a non-significant difference. **D.** As for C, but analysing wild type, *atxr5 atxr6*, *ligaseIV*, *mom1* and *smc4-1* mutants in a Col/Col homozygous background. **E.** As for C, but with genotypes in a Col/Ler hybrid background, and with the addition of *recq4a recq4b*. **F.** Comparison of crossover frequency (cM/Mb) in wild type, *recq4a recq4b* and *HEI10*, generated by SSLP mapping in Col/Ler F_2_ populations. The position of the *CTL3.9* T-DNAs are indicated by green and red vertical lines. The density of *CEN178* on the forward (red) and reverse (blue) strands are plotted. Crossover counts per interval were compared between genotypes using chi-square tests. Asterisks indicate intervals that had significantly different counts between wild type and *HEI10* or *recq4a recq4b*.

### Genetic control of crossover frequency within *CENTROMERE3*

To investigate genetic control of centromere-proximal crossovers we crossed *CTL3.9* to mutants with changed recombination or chromatin pathways, in inbred (Col/Col) backgrounds. The mutants used were, (i) *chromomethylase3* (*cmt3-11*), which is required for maintenance of CHG DNA methylation [59], (ii) *methyltransferase1* (*met1*), which is deficient in maintenance of CG DNA methylation [60], (iii) *atxr5 atxr6*, which are required for maintenance of the heterochromatic mark H3K27me1 [29], (iv) *smc4-1*, which disrupts a condensin complex required for gene silencing and chromosome compaction [61], (v) *mom1*, which is defective in DNA-methylation independent gene silencing and regulates transcription of *CEN178* repeats [62], (vi) *HEI10*, which over-expresses a pro-crossover E3 ligase [36], (vii) *ligase4*, which is required for non-homologous end joining (NHEJ) and DSB repair in repetitive regions [63], and (viii) *recq4a recq4b*, which is deficient in DNA helicases that negatively regulate crossover frequency [64].

We observed that *cmt3-11* caused a significant increase in *CTL3.9* crossover frequency (Wilcoxon test *P*=6.60⨉10^-4^) (**Fig. 5C** and **Table S3**), consistent with previous observations [44]. *CTL3.9* was crossed to generate *met1-3/+* heterozygous plants, which have reduced crossovers comparable to *met1-3* homozygotes yet have higher fertility [45]. *CTL3.9* crossover frequency was significantly reduced in *met1-3/+* (Wilcoxon test *P*=0.048) (**Fig. 5C** and **Table S3**). Hence, CG and CHG context DNA methylation maintenance have antagonistic effects on centromere-proximal crossover rate. No significant change in *CTL3.9* crossover frequency occurred in *smc4*, *atxr5 atxr6* or *lig4*, and a weak yet significant difference was observed in *mom1* (Wilcoxon test *P*=0.029) (**Fig. 5D** and **Table S8**).

In Arabidopsis, crossovers are generated by Class I and Class II pathways, which differ in their sensitivity to interference and interhomolog genetic polymorphism [52,53,65]. Therefore, we investigated *CTL3.9* crossovers in wild type, *cmt3*, *met1/+*, *HEI10* and *recq4 recq4b* Col/Ler F1 hybrids, to compare with the respective inbreds (**Fig. 5C-5E** and **Table S4**). Increased and decreased *CTL3.9* crossover frequencies were replicated in *cmt3* (Wilcoxon test *P*=5.80⨉10^-3^) and *met1/+* hybrids (Wilcoxon test *P*=1.12⨉10^-9^), showing that these changes are insensitive to interhomolog polymorphism (**Fig. 5E** and **Table S4**). The *recq4a recq4b* mutant causes increased Class II crossovers in the chromosome arms, whereas *HEI10* overexpression increases the Class I pathway [32,36,66]. Despite strong effects on recombination in the chromosome arms [32,64], *recq4a recq4b* did not significantly change *CTL3.9* crossovers, whereas *HEI10* showed a relatively weak but significant increase (Wilcoxon test *P*=5.55⨉10^-4^) (**Fig. 5E** and **Table S4**).

Although overall crossover rate was not increased, we performed further genetic mapping within *CTL3.9* to determine if spatial crossover patterns were changed in *HEI10* or *recq4a recq4b* hybrids. We genotyped 90 crossovers in wild type, 92 in *recq4a recq4b* and 90 in *HEI10* using ten Simple Sequence Length Polymorphism (SSLP) markers distributed evenly throughout *CTL3.9* (**Fig. 5F** and **Table S9**). In *recq4a recq4b*, crossover frequency significantly increased in the interval closest to the green T-DNA insertion, and decreased next to the red T-DNA, and at the left distal LRZ border (Chi square tests *P*=<0.05) (**Fig. 5F** and **Table S9**). In *HEI10*, a significant crossover change was observed close to the left distal LRZ border (Chi-square test *P*=<0.05) (**Fig. 5F** and **Table S9**). Although significant changes were observed in a minority of intervals, *HEI10* and *recq4a recq4b* had relatively limited effect on the *CTL3.9* recombination rate and landscape.

### The centromere-proximal recombination landscape is remodelled in *cmt3* and *met1/+* DNA methylation mutants

As DNA methylation defects changed *CTL3.9* crossover frequency in a hybrid background, we sought to perform high-resolution recombination mapping in *cmt3* and *met1/+*. Using the same fluorescent selection approach as for wild type, we mapped 1,033 crossovers within *CTL3.9* from *cmt3*, and 962 from *met1/+* (**Fig. 6A** and **Table S5**). The crossover landscape was significantly correlated between wild type, *cmt3* and *met1/+* (wt vs *cmt3 r=*0.743, wt vs *met1 r*=0.852) (**Fig. 6B-6D** and **Table S5**). To test for regional recombination changes within *CTL3.9*, we tallied crossovers in the NRZ, LRZs and distal regions and compared wild type, *cmt3* and *met1/+*. The LRZs showed significantly higher crossovers in *cmt3* (18.3%, chi test *P*=2.23x10^-5^), and significantly lower crossovers in *met1/+* (7.0%, chi test *P*=1.17x10^-3^), compared to wild type (11.4%) (**Fig. 6B-6D** and **Table S5**). In contrast, the distal regions significantly decreased in *cmt3* (81.6%, chi test *P*=2.82x10^-5^), and increased in *met1* (93.0%, chi test *P*=1.17x10^-3^), compared to wild type (88.6%) (**Fig. 6B-6D** and **Table S5**). Only a single NRZ crossover was observed in *cmt3*, indicating that the NRZ are stably repressed for crossover across these genotypes. This shows that relative crossover distributions have changed within *CTL3.9* between wild type, *cmt3* and *met1/+*.

**Figure 6.**
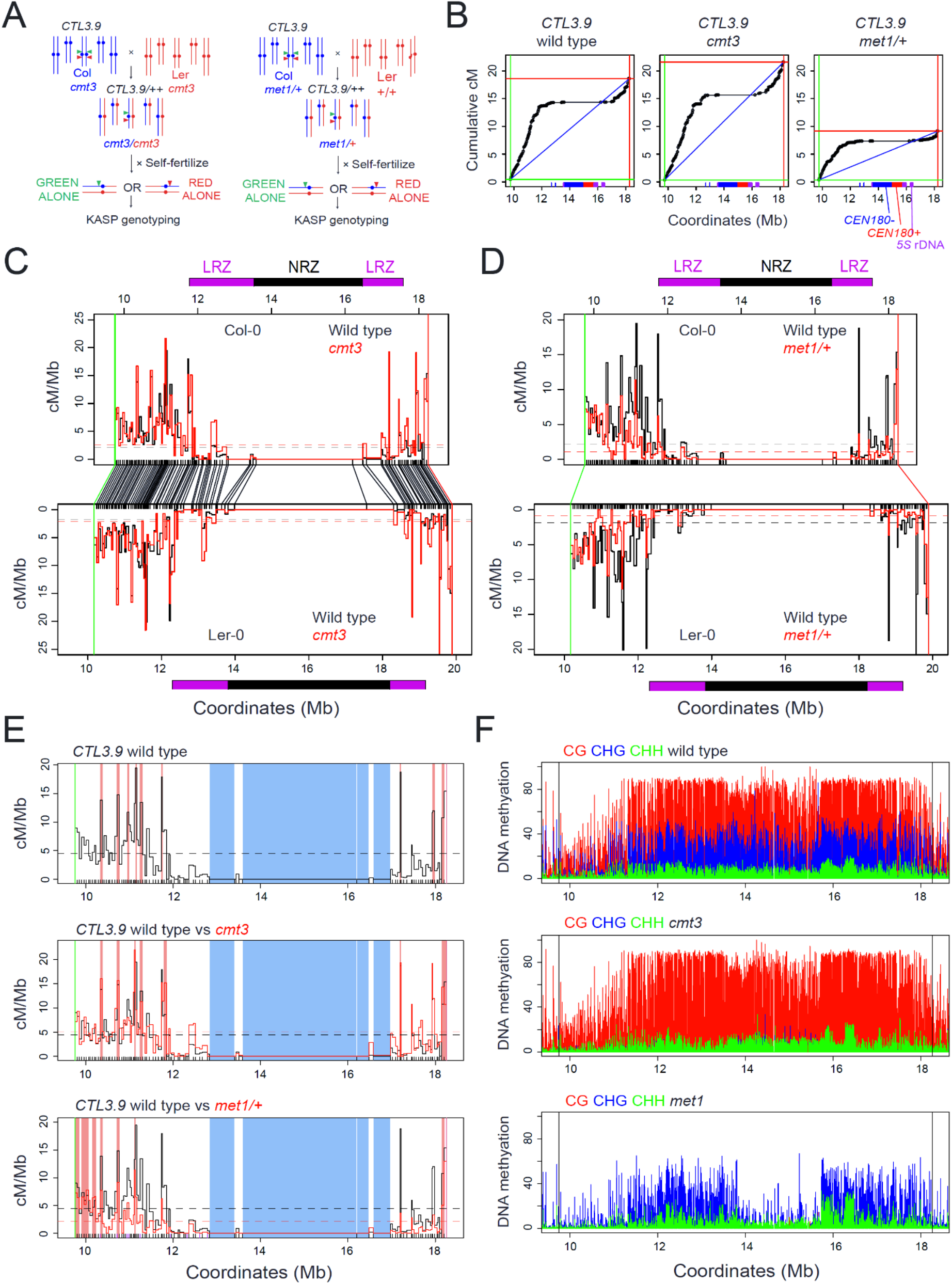
Remodelling of the centromere-proximal crossover landscape in wild type, *cmt3* and *met1/+*. **A.** Genetic strategy to recover crossovers within the *CTL3.9* centromeric FTL interval from *cmt3* and *met1/+* mutants. FTL T-DNAs encoding red and green fluorescent proteins are indicated by triangles. The parental chromosomes are from the Col (blue) and Ler (red) accessions. **B.** Cumulative genetic map (centiMorgans, cM) relative to *CTL3.9* genomic coordinates against the Col-CEN assembly in wild type, *cmt3* and *met1/+*. The blue diagonal lines show a linear relationship, with the red and green vertical/horizontal lines showing the location of the *CTL3.9* T-DNAs. The position of *CEN178* satellite repeats are shown as red (forward) and blue (reverse) ticks on the x-axis, in addition to *5S* rDNA (purple). **C.** Plots of crossover frequency (centiMorgans per Megabase, cM/Mb) within *CTL3.9* in wild type (black) and *cmt3* (red) projected against the Col (upper), or Ler (lower) assemblies, with mean values shown by dotted horizontal lines. The position of KASP genotyping markers are indicated as black x-axis ticks, and *CTL3.9* T-DNAs are indicated by red and green lines, and connected between the maps by lines. The LRZs (purple) and NRZ (black) as defined in wild type are shown as coloured blocks above and below the plots. **D.** As for C, but plotting wild type (black) and *met1/+* (red) crossover frequency (cM/Mb). **E.** *CTL3.9* crossover frequency (cM/Mb) in wild type (upper), *cmt3* (middle) and *met1/+* (lower), with significantly high (hot spots, pink) and low (cold spots, blue) intervals shown by coloured shading. **F.** Plots of DNA methylation (%) across *CTL3.9* derived from BS-seq data in wild type, *cmt3* and *met1* [87,88], coloured according to CG (red), CHG (blue) and CHH (green) sequence contexts. The position of the flanking *CTL3.9* T-DNAs are indicated by black vertical lines.

We tested for significantly hot or cold intervals in *met1/+* and *cmt3* compared to the random expectation. This identified that the large central *CS* cold spots maintained crossover suppression in both mutants (**Fig. 6E** and **Tables S5-S6**). The *cmt3* mutant showed fewer hotspots (*n*=8) than wild type (*n*=10), six of which were in the same locations (**Fig. 6E** and **Tables S5-S6**). In *met1/+*, the fine-scale landscape was more significantly changed compared to wild type, with hotspots in proximity to the *CEN178* arrays being strongly suppressed (**Fig. 6D-6E** and **Tables S5-S6**). For example, *HS6* and *HS7* are no longer significant hotspots in *met1/+*. Instead, new *met1/+* hotspots are detected close to the distal boundaries of the interval, with nine *met1/+* hotspots observed that were not present in wild type, together with five that were shared between genotypes (**Fig. 6E** and **Tables S5-S6**). Together this shows distinct patterns of remodelling of the crossover landscape within *CTL3.9* in *cmt3* and *met1/+*.

The changes in *cmt3* crossover distribution correlates with a dramatic reduction in CHG context DNA methylation within the centromere and pericentromere (**Fig. 6F**). Hence, loss of CHG methylation increases centromere-proximal crossover frequency, but the overall topology of the recombination landscape remains similar to wild type. The *met1* mutant shows a strong reduction in CG context DNA methylation across the centromere region, in addition to depletion of non-CG methylation within the *CEN178* arrays (**Fig. 6F**) [22]. These distinct changes to the *met1* centromeric DNA methylation landscape associate with a reduction and redistribution of crossovers, which was distinct from the crossover changes observed in *cmt3*. This demonstrates the contrasting effects of CG and CHG context DNA methylation on the centromere-proximal recombination landscape in Arabidopsis.

## Discussion

Despite playing a conserved function in kinetochore complex assembly, the size, DNA sequence and structure of eukaryotic centromeres are highly divergent within and between species [19–21]. Although centromeres are diverse at the DNA sequence level, they also share the feature of suppressed meiotic crossovers across species [8,10,12,22]. It has been proposed that suppression of centromere-proximal crossovers reduces gamete aneuploidy [8,14,67]. Consistently, we observed that crossovers are completely absent within the centromere NRZs, which contain the *CEN178* satellite arrays and regions of CENH3 enrichment, implying that NRZ crossovers may be deleterious in Arabidopsis. Additionally, we identified 1-2 Mb pericentromeric LRZs that surround the NRZs and experience strongly suppressed crossovers. The low levels of crossovers that occur within the LRZs are locally enriched with isolated euchromatic gene islands.

We propose that centromere crossover suppression is caused by a combination of NRZ structural genetic polymorphism and repressive heterochromatic marks, including dense DNA methylation and H3K9me2. We propose that centromere-proximal crossover suppression arises as, (i) the number of DSB precursors occur at lower levels [22,39,44], and that (ii) heterochromatin, and (iii) structural polymorphism, further inhibit downstream crossover repair pathways. A fourth possibility is that the centromere and kinetochore complex actively recruit factors that suppress crossover repair, or that the kinetochore might impose physical constraints that suppress crossover formation. As the centromeric and pericentromeric regions are enriched for REC8-cohesin [28], by analogy with budding yeast this may promote meiotic DSB repair using a sister chromatid, contributing to crossover suppression [8,68]. Meiotic DSBs could also be channelled into non-crossover repair [2], thereby limiting centromere-proximal crossovers. In comparison, human centromeres are composed of similar megabase-scale α-satellite arrays, although narrower (∼100 kb) regions of CENP-A enrichment are observed, which are DNA hypomethylated in the CG sequence context [23,25]. As human centromeres are also suppressed for meiotic crossovers [10], species-specific chromatin and DNA sequence organisation may play varying roles in shaping centromere proximal recombination landscapes.

We demonstrate that changes to the DNA methylation landscape, via mutation of either the CG (*met1/+*) or CHG (*cmt3*) maintenance pathways, cause contrasting changes to the landscape of centromeric crossover frequency. In wild type, a zone of crossover suppression extends from the NRZ into the flanking LRZs. In *met1/+* mutants, the zone of LRZ crossover suppression appears to extend and causes clustering of new recombination hotspots in more distal locations. This effect is reminiscent of the long-range effects of crossover interference [69], and we propose that a *cis*-acting recombination-suppressing signal emanating from the centromere may be strengthened in *met1*. In contrast, loss of CHG DNA methylation in *cmt3* increases LRZ crossovers, at the expense of the distal regions, although the NRZ remains strongly crossover-suppressed. Previous work revealed that both *met1* and *cmt3* experience greater levels of pericentromeric meiotic DSBs, measured by SPO11-1-oligo sequencing [39,44]. Hence, divergent changes occur to the centromere-proximal crossover landscape in these mutants, despite similar increases to the initiating DSBs [39,44]. This implies that DNA methylation context, and associated chromatin marks, have varying effects on meiotic recombination repair, downstream of SPO11-1-dependent DSBs. Further profiling of meiotic recombination, including single strand invasion, joint molecule formation and crossover resolution in these mutants may thus be revealing.

We observed a significant number of genes in the LRZs and NRZs that showed evidence of gene expression from RNA-seq data. Consistent with population genetics expectations for crossover-suppressed regions [70], these genes have significantly higher levels of genetic diversity than in the chromosome arms. As crossovers occur at low levels in the centromere proximal regions, mutations in these genes may not be efficiently purged via selection and therefore may accumulate over time, although other evolutionary forces, including mutation rates, may also contribute to the observed higher genetic diversity. LRZ/NRZ genes experience similar levels of SPO11-1-oligo formation in their promoters and terminators compared with genes located in the chromosome arms. As the LRZ/NRZ genes show elevated levels of the heterochromatic marks DNA methylation and H3K9me2, we propose that despite initiation of meiotic recombination, downstream crossover repair fate is inhibited. The LRZ and NRZ are also strongly enriched for Gypsy/Ty3 retrotransposons, which may reflect integration bias, or post-insertion effects of selection removing insertions in the chromosome arms. One consequence of gene and transposon co-location within the NRZ/LRZs is that they will tend to maintain linkage with each other and the centromere. The extent to which centromere proximal recombination suppression is important for chromosome segregation will be important to explore, in addition to how linked-inheritance influences genetic variation and evolution of genes and transposons resident in these regions.

## Methods

### Plant material and growth

*Arabidopsis thaliana recq4a-4* (GABI_203C07), *recq4b-2* (N511330/N660303) [71], *cmt3-11* (N648381) [72], *met1-3* (N16394) [60], *atxr5* (N630607), *atxr6* (N866134) [29], *mom1* (N826153) [62], *ligaseIV* (N656431) [63] and *smc4-1* (N69854) [61] mutants, the traffic line *CTL3.9* [48], and the *HEI10* over-expressor line ‘C2’ [36], were generated in the Col background. The *met1-3* allele was backcrossed into Ler [45], and the *cmt3-7* (N6365) and *recq4a-W387X* (EMS) [64] mutants are in the Ler accession [59]. Plants were grown in controlled environment chambers at 20°C under long day conditions (16/8 hours light/dark photoperiods) with 60% humidity and 150 µmol m^-2^ sec^-1^ light intensity. To stratify germination, once seeds were sown on commercial soil, they were kept for two days in the dark at 4°C before transferring them into growth chambers.

### Mapping crossovers from sequencing data and NRZ and LRZ identification

Whole-genome Illumina re-sequencing datasets of Arabidopsis Col (SRX202246) and Ler (SRX202247) were downloaded from the NCBI SRA database [73]. The paired-end raw reads were evaluated for quality using FastQC (version 0.11.9), and then adapter and low-quality sequences were trimmed using Trimmomatic (version 0.38) [74]. BWA (version 0.7.15-r1140) was used to align short reads with default parameters to the Col-CEN reference genome [22,75] (https://github.com/schatzlab/Col-CEN). Read alignments with a mapping quality greater than 20 were considered as uniquely mapped and used in subsequent analysis. Tandem repeat finder (version 4.09) was used with default parameters to scan the centromere-masked genome for tandem repeats, and used for quality examination of proximal SNPs [76].

To obtain high-confidence SNPs that can differentiate Col and Ler genotypes, we used the following strategy. First, SNPs and structural variants (SVs) were predicted from the sequencing datasets of Col and Ler accessions using inGAP-family [77]. Then, SNPs were filtered to remove potential false positives that arise from sequencing errors, small indels, tandem repeats and SVs using inGAP-family, as described previously [77,78]. Further, the SNPs were evaluated and filtered in both BC_1_ and F_2_ populations. Whole-genome Illumina resequencing datasets of Col/Ler BC_1_ (E-MTAB-11254) and F_2_ (E-MTAB-8165) populations were downloaded from the ArrayExpress database at EMBL-EBI, and aligned to Col-CEN, as above [9,33]. For the BC_1_ population data, SNPs were retained if; (i) the Col allelic ratio (Col read number divided by total read number) was larger than 0.6 and less than 0.9, and, (ii) the homozygous Col allelic ratio (number of samples with 0/0 genotype divided by the total number of genotyped samples) was larger than 0.4 and less than 0.7, and, (iii) the heterozygous Col allelic ratio (number of samples with 0/1 genotype divided by total number of genotyped samples) was larger than 0.3 and less than 0.6, and, (iv) the homozygous Ler allelic ratio (number of samples with 1/1 genotype divided by total number of genotyped samples) was less than 0.1, and, (v) the total number of genotyped samples was larger than 70 (5-quantiles). For the F_2_ population, SNPs were retained if, (i) the Col allelic ratio was larger than 0.3 and less than 0.7, and, (ii) the homozygous and heterozygous Col, and the homozygous Ler allelic ratios were between 0.1 and 0.9, and, (iii) the total number of genotyped samples was larger than 5. Following both BC_1_ and F_1_ filtering, a set of 334,680 high-quality Col/Ler SNPs were retained for subsequent analysis.

For each BC_1_ and F_2_ sample, the read count and genotype profile of the filtered high quality SNPs were produced by inGAP-family, and then a sliding window-based method was adopted for detecting crossovers, using a window of 70 kb, and a step size of 35 kb [33–35]. Final crossover positions were further refined by examining the genotype information of individual proximal-SNPs. Poorly covered (<0.1× depth) and potentially contaminated samples (>2% of windows with first-allele frequency in the range of 0.8 to 0.9), were removed from further analysis. To further filter false crossovers caused by mis-genotyping, we calculated the double-crossover frequency, defined as the number of samples with double-crossovers divided by the number of samples with crossovers in the given window, for every 1 Mb window, with a 500 kb step size. We filtered double-crossovers in the given window, when the double crossover frequency was greater than 4%. Moreover, we manually examined crossovers that were located close to each side of the centromere of each chromosome for the BC1 and F2 populations, respectively. To be retained, a qualified crossover must be supported by sufficient coverage of Col and Ler specific reads, with a minimum of five accumulated reads, in both homozygous and heterozygous genotypes. A total of 1,009, 978, and 12,410 crossovers were retained for the female BC1, male BC1, and F2 populations, respectively. Crossovers were then tallied in 10 kb windows along the Col-CEN genome. The non-recombining zones (NRZs) were defined as the contiguous regions containing the main *CEN178* arrays where crossovers were never observed. From the boundaries of the NRZs, we defined the LRZs, as the 1 cM flanking regions. The 14,397 mapped crossovers were also tallied in the *CTL3.9* KASP marker windows to correlate recombination rates between experiments.

### Genome analysis and annotation

The Col-CEN genome was used for analysis [21,22]. The Ler assembly is from ENA study ID PRJEB55353 and corresponds to ENA assembly ID GCA_946406525 [21]. Gene, transposon and tandem repeat annotation for the Col and Ler genome assemblies were as reported [21,22]. StainedGlass (version 0.5) was used to generate sequence identity heat maps, using a 10,000 bp window size, to compare the repeat architecture of Col and Ler centromeres [79]. Minimap2 (version 2.24) and SYRI (version 1.6) were used to map regions of synteny and inversions between the Col and Ler assemblies [40,80]. To map the *CTL3.9* recombination data onto the Ler assembly, the sequences surrounding the KASP SNPs were aligned to Ler 0 using LASTZ (version 1.04.15).

### Analysis of genes and transposons in the LRZs and NRZs

Using the LRZ and NRZ coordinates against the Col-CEN assembly, we identified contained genes. These genes were masked for those located within the mitochondrial genome insertions on chromosome 2, and those located in the large pericentric Col/Ler inversion on chromosome 4 [41]. Fine-scale profiles around genes located in the LRZs (n=542), NRZs (n=132), or the chromosome arms (n=27,499), or the same number of randomly positioned loci of the same number and width distribution within the same regions, were calculated for ChIP-seq, RNA-seq, and bisulfite-seq data sets by providing bigWig files to the computeMatrix tool from deepTools (version 3.1.3) in ‘scale-regions’ mode [81]. Each feature was divided into non-overlapping, proportionally scaled windows between start and end coordinates, and flanking regions were divided into 10 bp windows. Mean values for each data set were calculated within each window, generating a matrix of profiles in which each row represents a feature with flanking regions and each column a window. Coverage profiles for an input sequencing library and a gDNA library were used in conjunction with those for ChIP-seq and SPO11-1-oligo libraries, respectively, to calculate windowed log_2_([ChIP+1]/[control+1]) coverage ratios for each feature. Meta-profiles (windowed means and 95% confidence intervals) for each group of features were calculated and plotted using the feature profiles in R (version 4.0.0).

We tested crossovers for overlap with LRZ genes using permutation tests, which compared the observed number of gene-overlapping crossovers with the numbers of gene-overlapping randomly positioned LRZ loci, across 10,000 permuted sets of the same number and widths as the crossovers. An equivalent procedure was performed to test for overlap of LRZ crossovers with transposon annotation. The length (kb) of transposon annotation was also calculated in 100 kb windows and plotted along the Col-CEN assembly for the Gypsy/Ty3, Copia/Ty1 and LINE-L1 superfamilies.

To investigate genetic diversity in LRZ and NRZ genes, permutation analysis of pairwise diversity (Pi) in genes was performed. We downloaded the variant call format (VCF) and annotation of the 1,135 A. thaliana natural accessions from Phytozome (https://phytozome-next.jgi.doe.gov) [46]. We calculated pairwise diversity (or π) for TAIR10 gene models, allowing for 15% missing calls per site. We removed sites that overlapped transposable elements, rDNA, plastid sequences and simple repeats in TAIR10 from the VCF file. We required that at least half of the length of the gene had sequencing coverage among the 1,135 accessions, in order to produce a reliable π calculation per gene. We excluded genes that overlapped inversions between the Col-CEN and Ler genome assemblies, in addition to genes that overlapped the mitochondrial insertion on chromosome 2 from the analysis. After filtering 336 LRZ and 58 NRZ genes were retained for analysis. We calculated median π for this filtered gene set separately for the NRZ and LRZ genes, and compared this value to 1,000 permutations of median π for genes in the chromosome arms.

### Assembly of the *CTL3.9* FTL line genome and comparison with Col-CEN

Oxford Nanopore sequencing of homozygous *CTL3.9* plants was performed as reported [22]. Reads were trimmed of adapter sequences using Porechop (version 0.2.4), and filtered for length and accuracy using Filtlong (version 0.2.0) (--min_mean_q 90, --min_length 30000). The trimmed and filtered reads were assembled using Flye (version 2.7) [82]. Flye contigs were scaffolded and orientated using RagTag (version 2.0.1) using the reference genome Col CEN. To compare the *CTL3.9* and Col-CEN assemblies, sequence identity dotplots were performed using ReDOTable (https://www.bioinformatics.babraham.ac.uk/projects/redotable/). T-DNA borders were identified by using LASTZ to search for the T-DNA left border sequence. EDTA was used to annotate transposons in the *CTL3.9* assembly [83]. DNA methylation in the *CTL3.9* genome was mapped using the ONT reads and Deepsignal-plant, as described [22]. The CG17 and CR55 T-DNAs were identified using the primers as listed in Table S11. Based on alignments of the T-DNAs to the *CTL3.9* genome assembly, we observed two T-DNA insertions in tandem at CG17. In comparison to the Col-CEN assembly, 17 bp in the *CTL3.9* assembly is missing close to the CG17 T-DNA insertion site. Likewise, the CR55 T-DNA insertions are in tandem. Moreover, a 1,289 bp region on one side of the CR55 T-DNA is duplicated on both sides of the insertion, which includes one gene (At3g06765) that encodes a non-coding RNA.

### Crossover measurement using *CTL3.9* seed fluorescence

*CTL3.9* comprises genetically linked T-DNAs expressing red or green fluorescent proteins in the seed from the *NapA* promoter that flank centromere 3 [48], which were used to measure crossover frequency and map recombination events. For each sample, three seed images were acquired; (i) brightfield, (ii) UV through a dsRed filter, and (iii) UV through a GFP filter, using a Fluorescent Stereomicroscope (Leica M165FC) [53]. CellProfiler (version 2.1.1) image analysis software was used to identify seed boundaries in micrograph images, and to assign RFP and GFP fluorescence intensity values to each seed object [53,84]. In a *CTL3.9 RG/++* hemizygous line, when a single crossover occurs between the T-DNAs, they are inherited separately through meiosis, resulting in seed with red or green fluorescence alone. *CTL3.9* genetic distance can then be calculated using the formula;

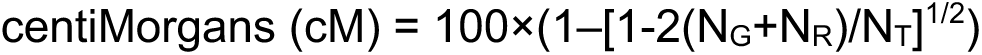

Where N_G_ is a number of green-alone fluorescent seeds, N_R_ is a number of red-alone fluorescent seeds, and N_T_ is the total number of seeds counted [53]. Statistical comparisons between samples were performed comparing the mean cM of replicate plants using Wilcoxon tests.

### *CTL3.9* KASP genotyping, crossover identification and analysis

Single nucleotide polymorphisms (SNPs) between the Col and Ler accessions and 50 base pairs of flanking sequence on each side of the SNP were used to design Kompetitive Allele Specific PCR (KASP) markers (LGC, Hoddesdon, UK). KASP uses two allele-specific forward primers and one common reverse primer. The two allele-specific primers posses unique tail sequences that correspond to a FRET (fluorescence resonant energy transfer) cassette; one labelled with FAM and the other with HEX. KASP allows differentiation of two alleles via competitive binding of allele-specific primers. If the genotype is homozygous, then only FAM or HEX fluorescent signals are observed. If the genotype is heterozygous, then a combination of FAM and HEX fluorescent signals is observed. The majority of SNPs analysed were located within genes, with the remainder located in intergenic regions.

F_2_ seeds showing green or red fluorescence alone were manually selected and grown on soil, alongside wild type and Col/Ler _F1_ heterozygote controls. Leaf tissue was collected when plants were three to four weeks old. LGC performed DNA extraction from leaf tissue and KASP genotyping. Raw fluorescence data for each marker were assessed for differentiation of genotypes and used to annotate markers in each sample as either Col/Col, Col/Ler or Ler/Ler genotype. Each plant was expected to contain a single crossover event within the *CTL3.9* interval, identified by a genotype transition from Col/Ler to Ler/Ler in green-alone seeds, or Ler/Ler to Col/Ler in red-alone seeds (**Fig. S6A**). The majority of plants showed an expected genotype transition associated with a single crossover (**Fig. S6A**).

A minority of plants (n=29) showed multiple genotype transitions, for example Col/Col to Col/Ler to Ler/Ler, or Ler/Ler to Col/Ler to Col/Col (**Fig. S6B**). These genotypes can be explained if homozygous red or green recombinant seeds were selected, in which case two crossovers on different chromatids are present (**Fig. S6B**). For these samples both independent crossovers were counted. Three plants were found to be entirely Col/Col genotype, which likely reflect seed contamination, and these samples were removed from analysis (**Fig. S6C**). Two remaining plants showed genotype transitions including Col/Ler to Col/Col to Col/Ler to Ler/Ler, or Ler/Ler to Col/Ler to Col/Col to Col/Ler, which are consistent with one recombinant chromatid and another chromatid with a double crossover event carrying an introgression from Col (**Fig. S6D**). In these cases, only one crossover chromatid was counted.

26 plants showed a missing genotype at the crossover site, meaning the recombination event could not be unambiguously placed in one of two adjacent intervals. In these cases the single crossover value was divided between the two marker intervals, in proportion to the unambiguous crossovers mapped within the same two intervals. For example, if interval A contained 11 crossovers, and interval B contained 9, then a value of 0.55 would be added to interval A, and 0.45 would be added to interval B, in order to place the ambiguous crossover. To define crossover hotspot and coldspot intervals, we calculated the expected number of crossovers per interval, assuming an even distribution, and compared this to observed events. Observed and expected crossover counts for each interval were used to perform chi-square tests, followed by Bonferroni correction for multiple-testing, to identify intervals that contained significantly higher or lower crossovers. This process was repeated separately for wild type, *met1/+* and *cmt3*.

### Simple Sequence Length Polymorphism (SSLP) marker design

To design SSLP genetic markers, chromosome 3 genomic DNA sequence from TAIR10 (Col) (GenBank CP002686.1) and Ath.Ler-0.MPIPZ.v1.0 (Ler) (GenBank LR215054.1) were used [31,85]. Mauve sequence alignment was performed between the Col and Ler sequences using Geneious Prime software. Primers were designed flanking identified Col/Ler indel polymorphisms. BLAST alignment to all chromosomes of the Col and Ler assemblies was performed on primer sequences to assess their uniqueness within the genome. Genetic markers were validated using leaf genomic DNA extracted from Col, Ler and F_1_ (Col/Ler) genotypes, in addition to no template controls.

### Profiling DNA methylation in Ler using Oxford Nanopore sequencing

This was performed, as reported [21,22]. Briefly, for genomic DNA extraction for Oxford Nanopore Technologies (ONT) sequencing, 3 week old *A. thaliana* plants, grown on 1⁄2 MS media containing 1% sucrose, were placed in the dark for 48 hours prior to harvesting. Approximately 10 grams of tissue were used per 200 ml of MPD-Based Extraction Buffer pH 6.0 (MEB). Tissue was flash frozen and then ground in liquid nitrogen using a pestle and mortar, and resuspended in 200 ml MEB. Ground tissue was thawed in MEB with frequent stirring. The homogenate was forced through 4 layers of Miracloth, and then filtered again through 4 layers of fresh Miracloth by gravity. Triton x-100 was added to a final concentration of 0.5% on ice, followed by incubation with agitation on ice for 30 minutes. The suspension was centrifuged at 800 *g* at 4°C for 20 minutes. The supernatant was removed and the pellet resuspended using a paintbrush in 10 ml 2-methyl-2,4 pentanediol buffer pH 7.0 (MPDB). The suspension was centrifuged at 650 *g* at 4°C for 20 minutes. The supernatant was removed and the pellet was washed with 10 ml of MPDB. Washing and centrifugation was repeated until the pellet appeared white, when it was resuspended in a minimal volume of MPDB. From this point onwards, all transfers were performed using wide-bore pipette tips. Five ml CTAB buffer was added to the nuclei pellet and mixed via gentle inversion, followed by incubation at 60°C until full lysis had occurred, taking between 30 minutes and 2 hours. An equal volume of chloroform was added and incubated on a rocking platform, with a speed of 18 cycles per minute, for 30 minutes, followed by centrifugation at 3000 g for 10 minutes. An equal volume of phenol/chloroform/isoamyl alcohol (PCI, 25:24:1) was added to the lysate, followed by incubation on a rocking platform (18 cycles per minute) for 30 minutes. The lysate was centrifuged at 3,000 *g* for 10 minutes and the upper aqueous phase was transferred into a fresh tube. The PCI extraction was then repeated. The extraction was repeated using only chloroform. One-tenth volume of 3M Sodium Acetate was added to the lysate and mixed by gentle inversion. Two volumes of ice cold ethanol were added and mixed by inversion. DNA was precipitated at -20°C for 48 hours. The precipitated DNA was removed using a glass hook and washed three times in 70% ethanol. The DNA was dissolved in 120 μl of 10 mM Tris-HCl pH 8.5.

Approximately 5 μg of DNA was size selected (>30 kb) using the BluePippin System (Sage Science) and the 0.75% agarose gel cassette (BLF7510, Biozym), using Range mode and BP start set at 30 kbp. Library preparation followed the ONT SQK-LSK109 protocol kit, using 1.2-1.5 μg of size-selected DNA in a volume of 48 μl. DNA was nick-repaired and end-prepped by the addition of 3.5 μl of NEBNext FFPE Buffer and NEBNext Ultra II End Prep Reaction Buffer, followed by 2 μl of NEBNext DNA Repair Mix and 3 μl NEBNext Ultra II End Prep Enzyme Mix (New England Biolab, E7180S), with incubation for 30 minutes at 20°C, followed by 30 minutes at 65°C. The sample was cleaned using 1×volume AMPure XP beads and eluted in 61 μl of nuclease-free water. Adapters were ligated at room temperature using 25 μl Ligation Buffer, 10 μl NEBNext T4 DNA Ligase and 5 μl Adapter Mix for 2 hours. The library was cleaned with 0.4×volume AMPure XP beads, washed using ONT Long Fragment buffer and eluted in 15 μl elution buffer.

We quantified CG, CHG and CHH context DNA methylation with DeepSignal-plant (version 0.1.4), which uses a deep-learning method based on a bidirectional recurrent neural network (BRNN) with long short-term memory (LSTM) units to detect 5mC methylation. R9 reads were filtered for length and accuracy using Filtlong (Version 0.2.0) (--min_mean_q 90, --min_length 20000). Base-called read sequence was annotated onto corresponding .fast5 files, and re squiggled using Tombo (version 1.5.1). Methylation prediction for the CG, CHG, and CHH contexts were called using DeepSignal-plant using the model: model.dp2.CNN.arabnrice2-1_120m_R9.4plus_tem.bn13_sn16.both_bilstm.epoch6.ckpt. The scripts ‘call_modification_frequency.py’ and ‘split_freq_file_by_5mC_motif.py’ provided in the DeepSignal-plant package were used to generate the methylation frequency at each CG, CHG and CHH site.

### Profiling CENH3 in Ler-0 using ChIP-seq

This was performed, as reported [21,22]. Briefly, approximately 12 g of 2-week old seedlings were ground in liquid nitrogen. Nuclei were isolated in nuclei isolation buffer (1 M sucrose, 60 mM HEPES pH 8.0, 0.6% Triton X-100, 5 mM KCl, 5 mM MgCl2, 5 mM EDTA, 0.4 mM PMSF, 1 mM pepstatin-A and 1×protease inhibitor cocktail), and crosslinked in 1% formaldehyde at room temperature for 25 minutes. The crosslinking reaction was quenched with 125 mM glycine and incubated at room temperature for a further 25 minutes. The nuclei were purified from cellular debris via two rounds of filtration through one layer of Miracloth and centrifuged at 2,500 g at 4°C for 25 minutes. The nuclei pellet was resuspended in EB2 buffer (0.25 M sucrose, 1% Triton X-100, 10 mM Tris-HCl pH 8.0, 10 mM MgCl2, 1 mM EDTA, 5 mM DTT, 0.1 mM PMSF, 1 mM pepstatin-A and 1×protease inhibitor cocktail) and centrifuged at 14,000 g at 4°C for 10 minutes. The nuclei pellet was resuspended in lysis buffer (50 mM Tris-HCl pH 8.0, 1% SDS, 10 mM EDTA, 0.1 mM PMSF and 1 mM pepstatin-A) and chromatin was sonicated using a Covaris E220 Evolution device with the following settings: power=150 V, bursts per cycle=200, duty factor=20% and time=90 seconds. Sonicated chromatin was centrifuged at 14,000 *g* and the supernatant was extracted and diluted with 1×volume of ChIP dilution buffer (1.1% Triton X-100, 20 mM Tris-HCl pH 8.0, 167 mM NaCl, 1.1 mM EDTA, 1 mM pepstatin-A and 1×protease inhibitor cocktail). The chromatin was incubated overnight at 4°C with 50 μl Protein A magnetic beads (Dynabeads, Thermo Fisher) pre-bound with 2.5 μl α-CENH3 (gift of Prof. Steven Henikoff). The beads were collected on a magnetic rack and washed twice with low-salt wash buffer (150 mM NaCl, 0.1% SDS, 1% Triton X-100, 20 mM Tris-HCl pH 8.0, 2 mM EDTA, 0.4 mM PMSF, 1 mM pepstatin-A and 1×protease inhibitor cocktail) and twice with high-salt wash buffer (500 mM NaCl, 0.1% SDS, 1% Triton X-100, 20 mM Tris-HCl pH 8.0, 2 mM EDTA, 0.4 mM PMSF, 1 mM pepstatin-A and 1×protease inhibitor cocktail). Immunoprecipitated DNA–protein complexes were eluted from the beads (1% SDS and 0.1 M NaHCO_3_) at 65°C for 15 minutes. Samples were reverse crosslinked by incubating with 0.24 M NaCl at 65°C overnight. Proteins and RNA were digested with Proteinase K treatment, and RNase A, and DNA was purified by phenol:chloroform:isoamyl alcohol (25:24:1) extraction and ethanol precipitation. Library preparation followed the Tecan Ovation Ultralow System V2 library protocol. ChIP samples were PCR amplified for 12 cycles and sequenced with 150 bp paired-end reads on an Illumina instrument by Novogene.

Deduplicated paired-end CENH3 ChIP-seq Illumina reads (2×150 bp) from Col and, Ler were processed with Cutadapt (version 1.18) to remove adapter sequences and low-quality bases (Phred+33-scaled quality <20). For each accession, trimmed reads were aligned to the respective genome assembly using Bowtie2 (version 2.3.4.3), using the following settings: -- very-sensitive --no-mixed --no-discordant -k 10 --maxins 500. Up to 10 valid alignments were reported for each read pair. Read pairs with Bowtie2-assigned MAPQ <10 were discarded using Samtools (version 1.10). For retained read pairs that aligned to multiple locations, with varying alignment scores, the best alignment was selected. Alignments with more than 2 mismatches or consisting of only one read in a pair were discarded. For each data set, bins per-million-mapped-reads (BPM; equivalent to transcripts-per-million, TPM, for RNA-seq data) coverage values were generated in bigWig and bedGraph formats with the ‘bamCoverage’ tool from deepTools (version 3.5.0). Reads that aligned to chloroplast or mitochondrial DNA were excluded from coverage normalisation.

## Acknowledgments

This work was supported by BBSRC grants BB/S006842/1, BB/S020012/1 and BB/V003984/1, European Research Council Consolidator Award ERC-2015-CoG-681987, Marie Curie International Training Network ‘MEICOM’ and Human Frontier Science Program award RGP0025/2021 to IRH, by core funding from the Max Planck Society to RM and DW, Alexander von Humboldt Foundation Fellowships to QL and JBF, EMBO Long-Term postdoctoral fellowships to JBF (ALTF 329-2018) and RB (ALTF 224-2022), a Human Frontiers Science Program (HFSP) Long-Term Fellowship (LT000819/2018-L) to FAR, ERA CAPS 1001G+ grant to DW, and a Broodbank Fellowship to MN. We thank Steven Jacobsen (UCLA), Scott Poethig (University of Pennsylvania) and Craig Pikaard (Indiana University) for providing seed stocks, and Adrian Gonzalo for scientific discussions.

## Author Contributions

Conceived and designed the analysis; JBF, MN, QL, RB, AJT, FAR, PW, DW, RM and IRH. Collected the data; JBF, MN, FAR, AH and REN.

Contributed data or analysis tools; JBF, MN, QL, RB, AJT, FAR, PW, DW, RM and IRH. Performed the analysis; JBF, MN, QL, RB, AJT, FAR, PW, DW, RM and IRH.

Wrote the paper. JBF, MN, QL, RB, AJT, FAR, PW, DW, RM and IRH.

## Ethics approval

Does not apply to this study.

## Supplemental Information

**Figure S1.**
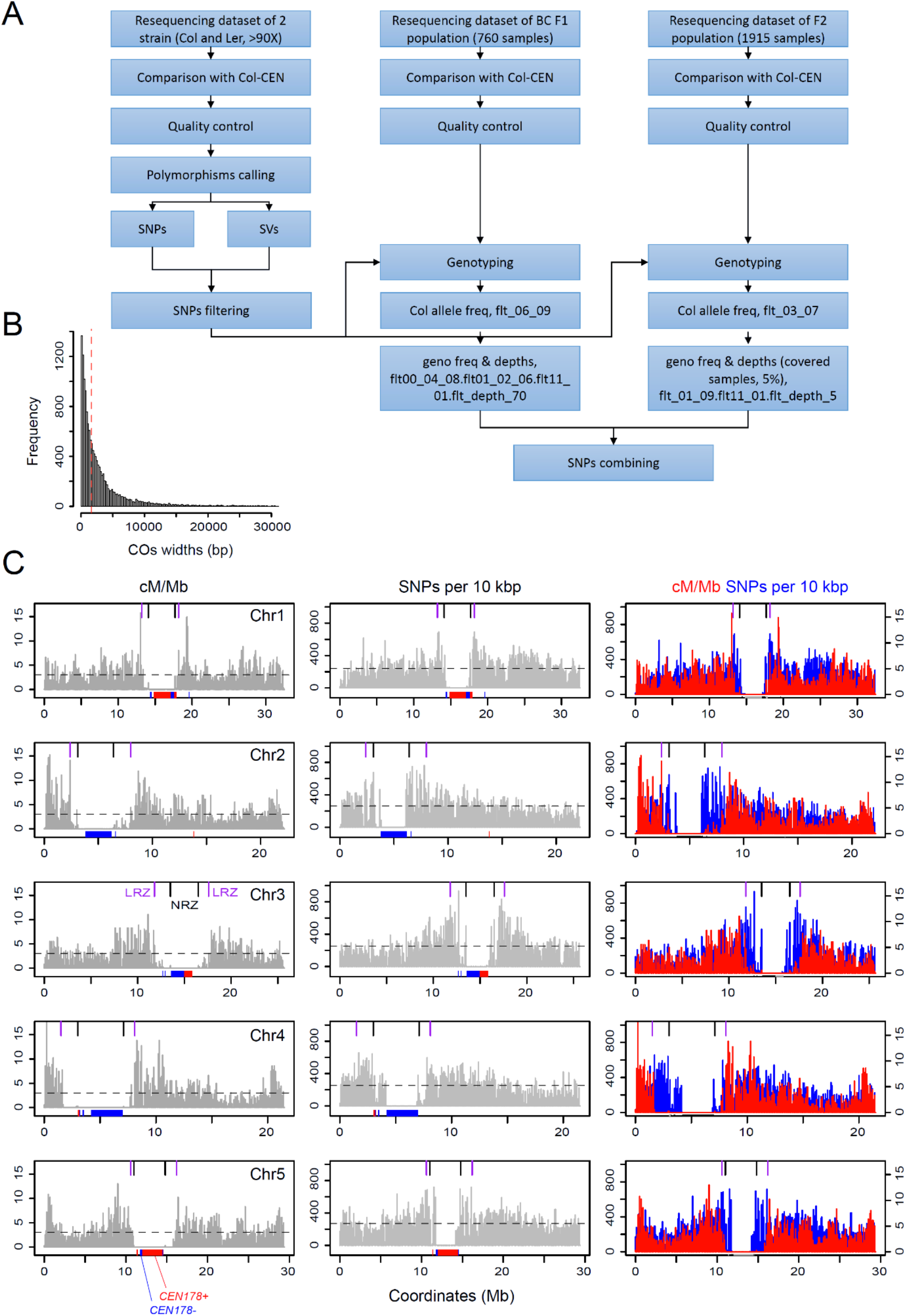
A refined pipeline for mapping crossovers from Col/Ler recombinant sequencing data. **A.** Schematic diagram showing the computational pipeline used to filter Col/Ler polymorphisms for SNPs that can be reliably used for crossover identification from backcross and F_2_ populations. **B.** Histogram of crossover widths (bp) identified by our mapping pipeline, with the mean value shown by the dotted red line. **C.** 10 kb windows are plotted along the Col-CEN assembly showing crossover frequency (cM/Mb, left), SNPs (middle) and an overlay (cM/Mb=red, and SNPs=blue) along each chromosome. The horizontal dotted lines indicate genome average values. NRZ (black) and LRZ (purple) boundaries, as shown in Fig. 1, are indicated as ticks along the upper axis. *CEN178* positions are indicated as ticks along the x axis (red=forward strand, blue=reverse strand).

**Figure S2.**
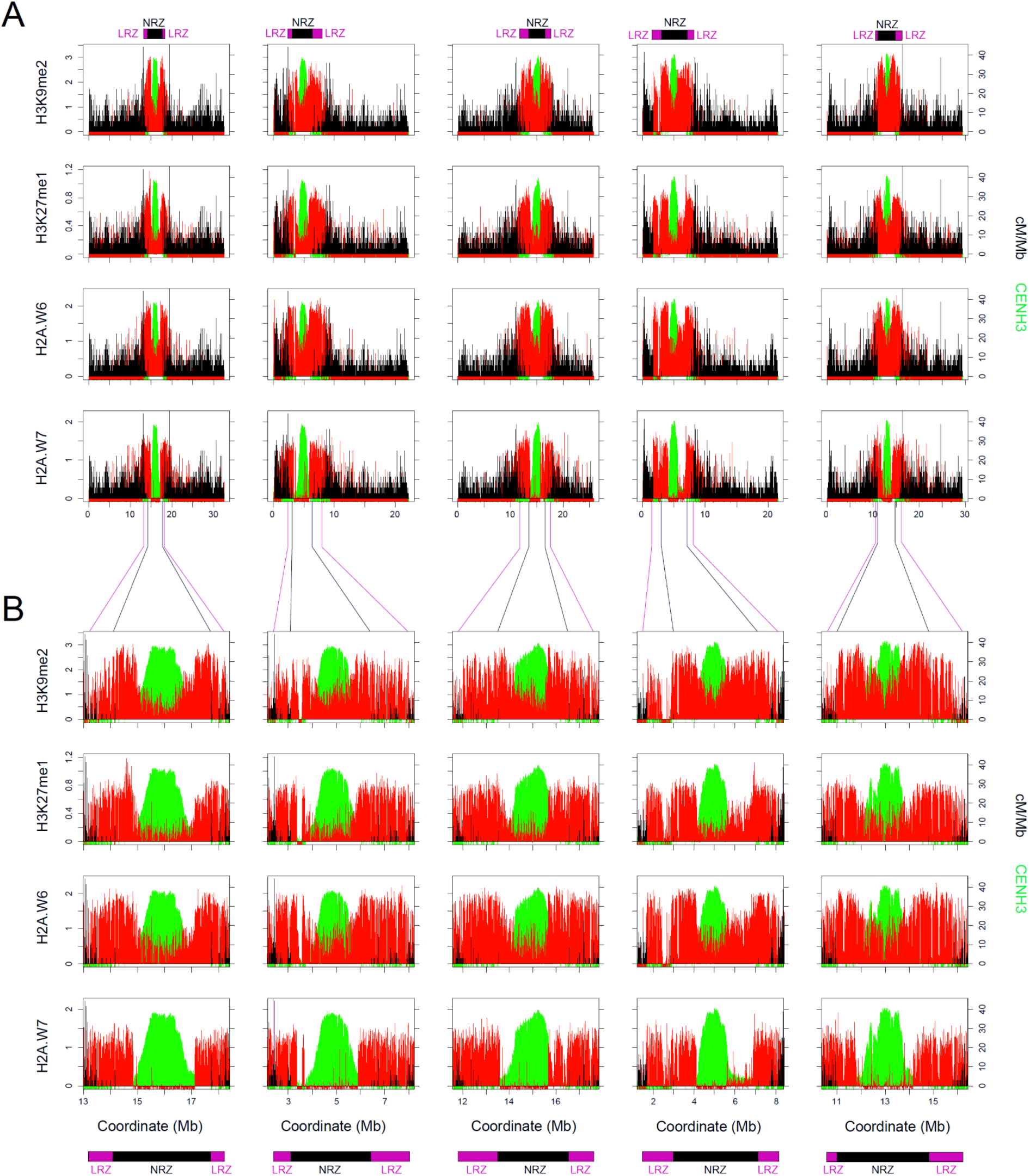
Zones of centromeric crossover suppression and heterochromatic histone modifications. **A.** Col/Ler crossover frequency (cM/Mb) mapped against the Col-CEN genome assembly in 10 kb windows is plotted (black). CENH3 ChIP-seq enrichment (green) is plotted using the same 10 kb windows. Above each plot, the location of the non-recombining zone (NRZ, black), and low-recombining zones (LRZ, purple), are indicated. Plots are shown compared to H3K9me2, H3K27me1, H2A.W6 and H2A.W7 ChIP-seq (red) enrichment for the same windows [28,29,37]. **B.** As for A, but showing a zoom of the NRZ and LRZ regions and the NRZ-LRZ positions are shown beneath by the black/purple bars.

**Figure S3.**
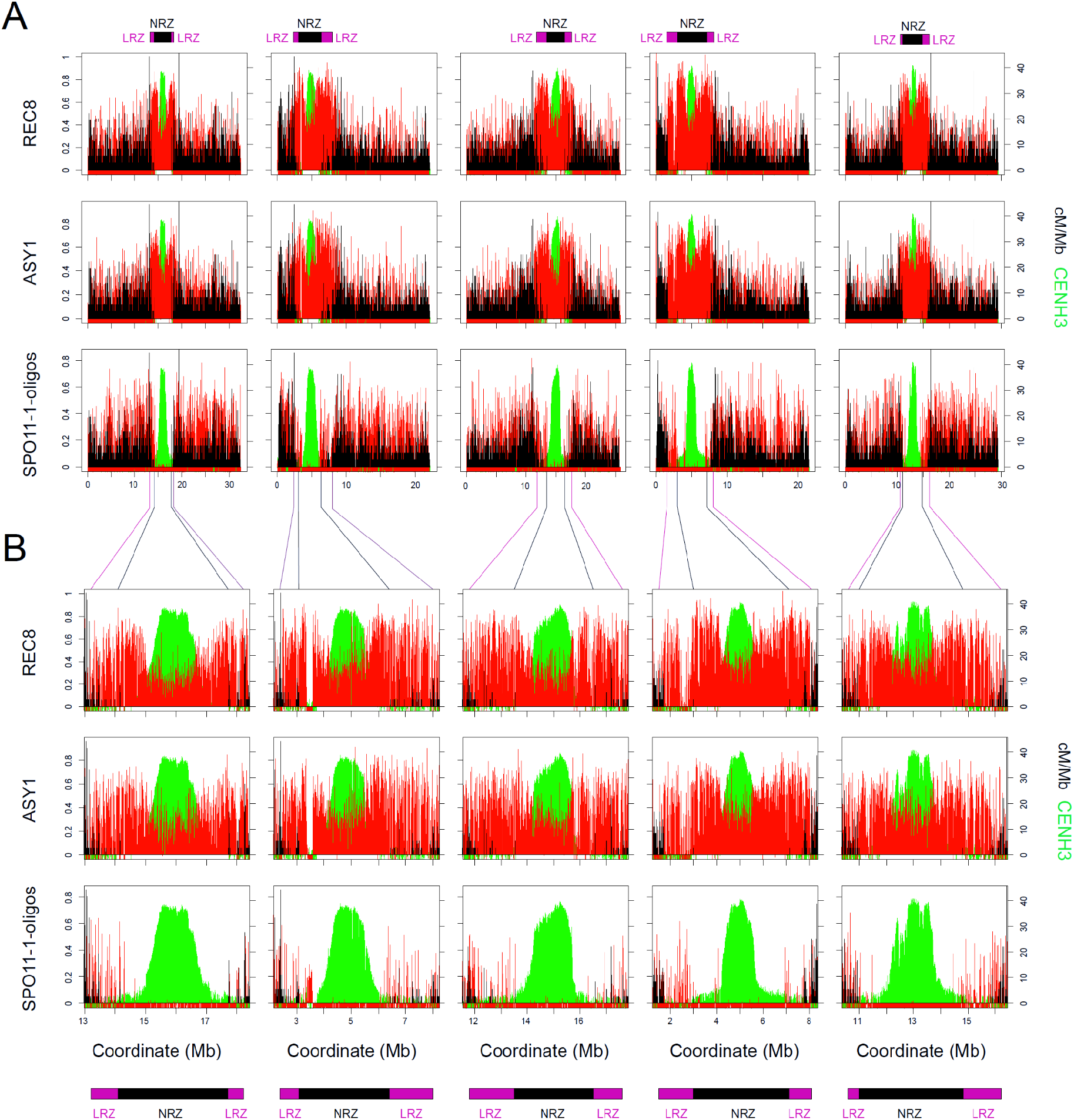
Zones of centromeric crossover suppression and REC8, ASY1 and SPO11-1-oligos. **A.** Col/Ler crossover frequency (cM/Mb) mapped against the Col-CEN genome assembly in 10 kb windows is plotted (black). CENH3 ChIP-seq enrichment (green) is plotted using the same 10 kb windows. Above each plot, the location of the non-recombining zone (NRZ, black), and low-recombining zones (LRZ, purple), are indicated. Plots are shown compared to REC8 and ASY1 ChIP-seq (red) enrichment, and SPO11-1-oligo enrichment, for the same windows [28,38,39]. **B.** As for A, but showing a zoom of the NRZ and LRZ regions and the NRZ-LRZ positions are shown beneath by the black/purple bars.

**Figure S4.**
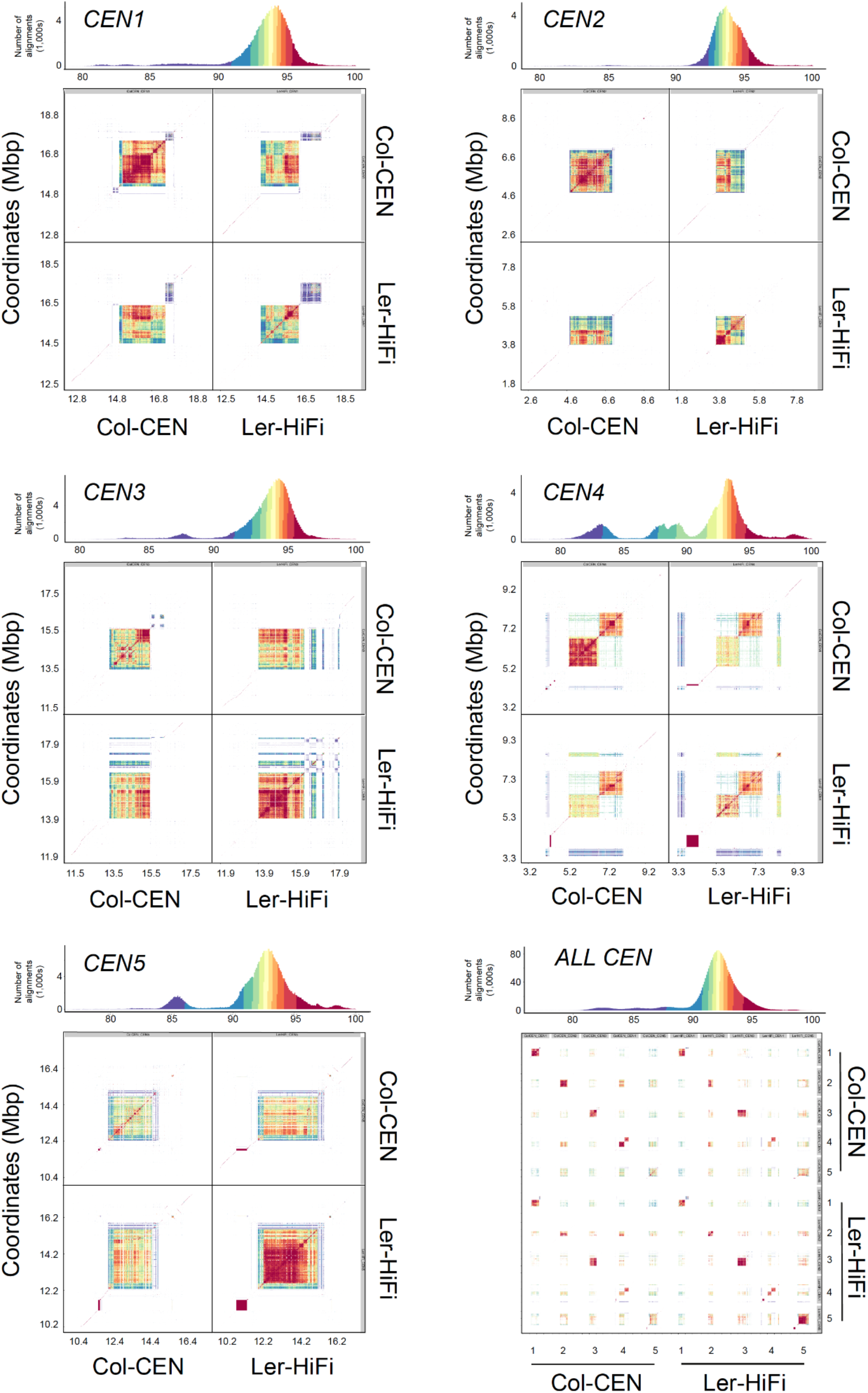
Structural comparison of Col and Ler *CEN178* centromere satellite arrays. Sequence regions including and surrounding the main *CEN178* satellite arrays were compared between the Col and Ler assemblies, for each chromosome, using sequence identity heat maps generated by StainedGlass [79]. In addition, a comparison of all centromeres between Col and Ler is shown (lower right). A histogram of % sequence identity values within each set of heat maps is shown above, which indicates color correspondences. Regions shaded red show highest levels of pairwise sequence identity.

**Figure S5.**
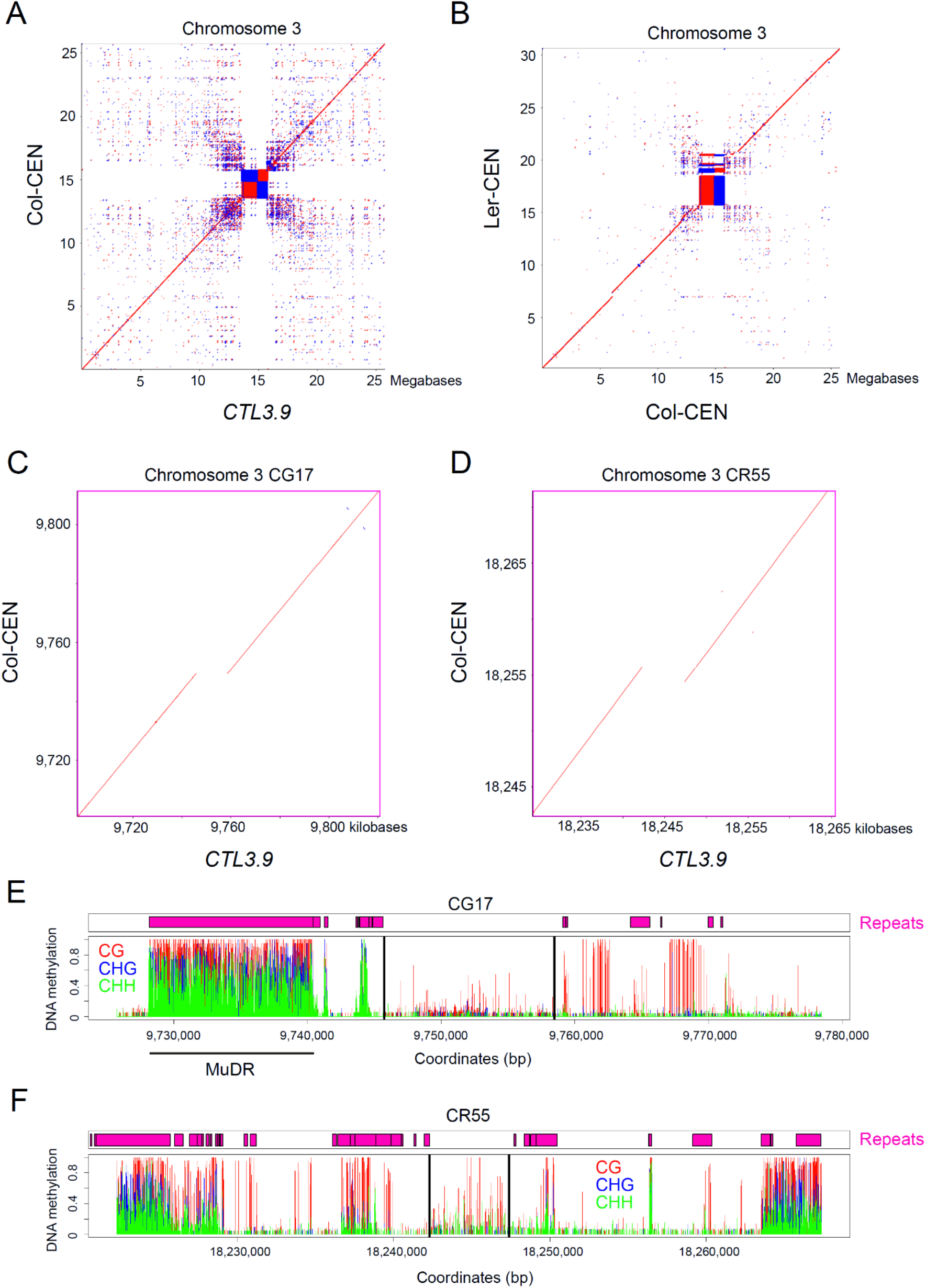
Genetic and epigenetic structure of *CTL3.9* fluorescent crossover reporter T-DNAs. **A.** Dot plot analysis of chromosome 3 from the Col (Col-CEN) and *CTL3.9* genome assemblies. Red and blue shading indicate forward and reverse strand similarity, respectively. **B.** As for A, but comparing chromosome 3 from the Col-CEN and Ler assemblies. **C.** As for A, but showing the region surrounding the CG17 *CTL3.9* T-DNA insertion. **D.** As for A, but showing the region surrounding the CR55 *CTL3.9* T-DNA insertion. **E.** ONT-based DNA methylation maps (proportion) for CG (red), CHG (blue) and CHH (green) sequence contexts across the CG17 T-DNA region. The T-DNA boundaries are indicated by vertical black lines. Above the plot, the pink rectangles indicate EDTA repetitive sequence annotation. **F.** As for E, but showing the region surrounding the CR55 T-DNA.

**Figure S6.**
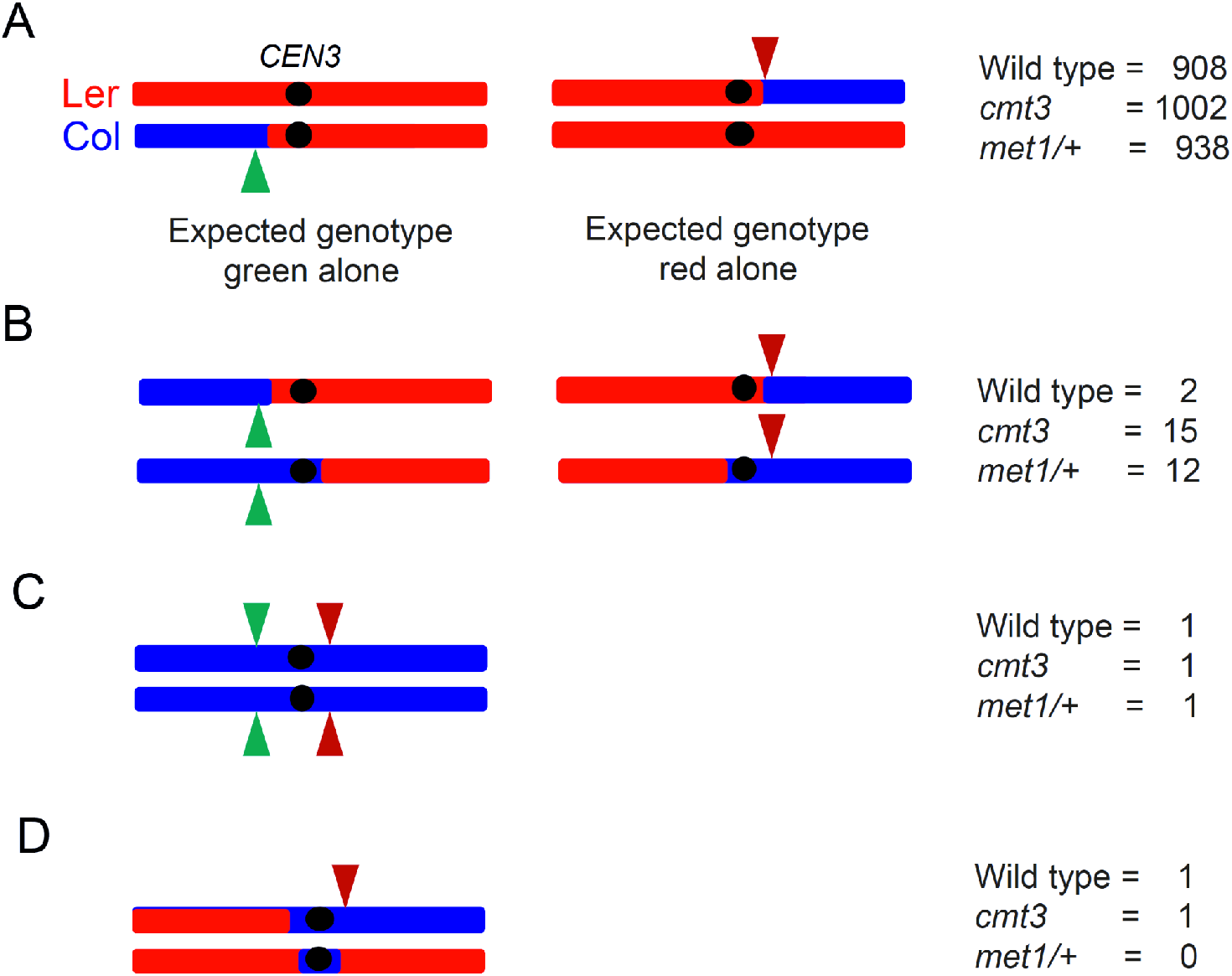
Genetic outcomes during *CTL3.9* fluorescent seed selection and genotyping. **A.** Diagrams showing the expected genotype of the majority of the seed selected as having green- or red-alone fluorescence, from a Col/Ler *CTL3.9/++* hybrid. Red and blue indicate Ler and Col genotypes, respectively. The position of centromere 3 (*CEN3*) is indicated with a black circle. The number of plants showing this genotype pattern for each population are shown to the right. **B.** In a small number of cases, the selected seed were homozygous red, or green, fluorescent, which can be explained by formation of the F_2_ individual from fertilisation of two independent crossover gametes. In these cases both crossover positions were retained for analysis. **C.** In these cases all genotypes were Col homozygous and are most likely explained by seed contamination, and so these samples were removed from analysis. **D.** Two samples showed an unexpected recombinant profile, consistent with three crossover events. This can be interpreted as one chromatid containing a single crossover producing a red-alone fluorescent phenotype. The other chromatid appears to have experienced a double crossover that resulted in a Col centromere introgression into an otherwise Ler background.

**Figure S7.**
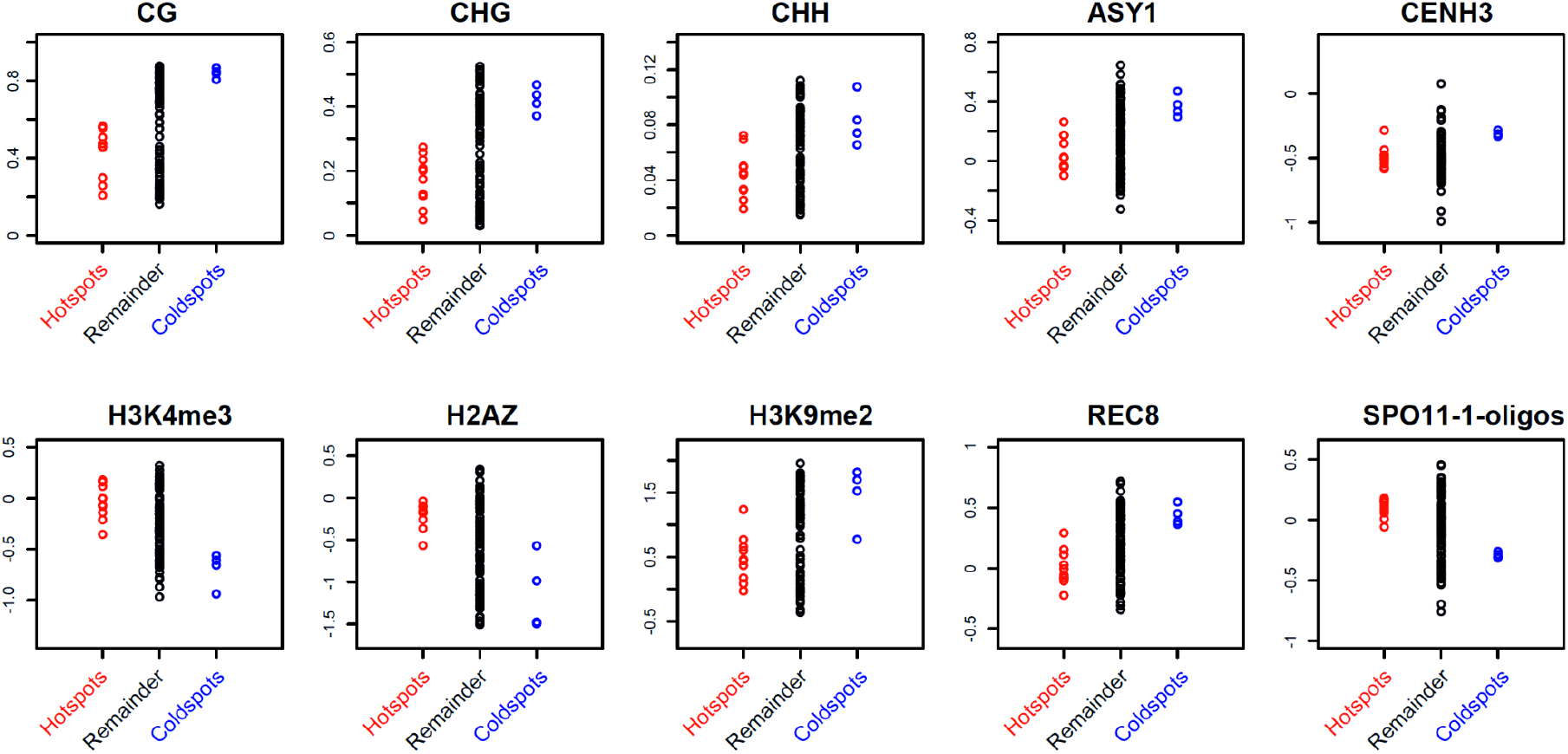
Chromatin and recombination states within *CTL3.9* hotspots and coldspots. *CTL3.9* map intervals that had significantly higher (hotspots, red) or lower (coldspots, blue) crossover frequency were analysed for multiple chromatin and recombination datasets and compared with the remainder (black) of the map intervals. Intervals are compared for ONT based DNA methylation levels in CG, CHG and CHH sequence contexts, ASY1, CENH3, H3K4me3, H2A.Z, H3K9me2, REC8 and SPO11-1-oligos [22,28,29,37–39].

**Table S1. Low-Recombining Zone and Non-Recombining Zone coordinates in the Col CEN assembly.** With reference to the Col-CEN assembly, the chromosome length (bp) is given, in addition to the left and right low recombining zone (LRZ) and non-recombining zone (NRZ) boundaries. The start and end of the main *CEN178* satellite arrays are also given, for each chromosome.

**Table S2. Genes located within the LRZs and NRZs.** The table provides whether genes are located in the LRZ or NRZ, their chromosome and start and end coordinate against Col-CEN, strand, gene name, TPM values for RNA-seq data generated from leaf or meiocytes [43], and gene annotation information from TAIR.

**Table S3. *CTL3.9* fluorescent data and crossover frequency measurements in Col/Col inbred backgrounds.** Crossover frequency measurements derived from fluorescent *CTL3.9/++* seed in wild type, *cmt3, met1/+* and *HEI10* overexpression lines. The ‘ctrl_1’ samples are the control that was used for comparison with *met1/+* and *HEI10* overexpression lines, while ‘ctrl_2’ were wild type controls for *cmt3*.

**Table S4. *CTL3.9* fluorescent data and crossover frequency measurements in Col/Ler hybrid backgrounds.** Crossover frequency measurements derived from fluorescent *CTL3.9/++* seed in wild type, *cmt3, met1/+, HEI10* overexpression and *recq4a recq4b* lines. The ‘ctrl_1’ is the control that was used for *recq4a recq4b*, and ‘ctrl_2’ for *cmt3*, and ‘ctrl_3’ for *met1/+* and *HEI10* overexpression lines.

**Table S5. *CTL3.9* crossover frequency maps in wild type, *met1/+* and *cmt3*.** For each genetic interval with the *CTL3.9* map, the number of crossovers observed are shown for wild type, *cmt3* and *met1/+*, in addition to cM/Mb values.

**Table S6. Crossover hot spots and cold spots observed in the wild type, *cmt3* and *met1/+ CTL3.9* recombination maps.** The table lists for the wild type, *cmt3* and *met1/+ CTL3.9* recombination maps, intervals that were significantly hot (hot spot, *HS*), or cold (cold spot, *CS*), based on observed and expected crossovers, assuming an even distribution. Observed and expected crossover counts for each interval were used to perform chi-square tests, followed by Bonferroni correction for multiple-testing, with the adjusted *P* value provided. Also provided are interval start coordinates and widths against the Col-CEN assembly, and the measured crossover rate (centiMorgans per Megabase, cM/Mb).

**Table S7. Crossover frequency map in the *HS6* hotspot.** 23 plants had a crossover event within the *HS6* hotspot. Recombinant plants were genotyped using four Col/Ler dCAPS markers (H6-1, H6-2, H6-3 and H6-4) to fine-map crossover locations.

**Table S8. *CTL3.9* fluorescent data and crossover frequency measurements in wild type and *smc4, atxr5 atxr6, mom1, ligase IV* mutant backgrounds**. Crossover frequency measurements derived from fluorescent *CTL3.9/++* seed in wild type, *smc4, atxr5 atxr6, mom1* and *ligaseIV* mutants. The ‘ctrl_1’ samples are wild type controls for *smc4*, ‘ctrl_2’ samples were wild type controls for *atxr5 atxr6*, ‘ctrl_3’ samples were wild types for *mom1*, and ‘ctrl_4’ samples were wild type controls for *ligase IV*.

**Table S9. *CTL3.9* SSLP genetic mapping in wild type, *recq4a recq4b* and *HEI10* overexpression lines.** Crossover numbers, and cM/Mb values, identified within each interval using Col/Ler SSLP genotyping are shown in wild type, *recq4a recq4b* and *HEI10* overexpression.

**Table S10. Col/Ler SNPs used as markers for KASP genotyping within the *CTL3.9*.** For each SNP, an id is assigned based on the TAIR10 coordinates of the SNP. IUPAC codes for the SNPs are indicated and the surrounding 50 bp region on either side of the SNP is shown. This data was provided to LGC (Hoddesdon, UK) to design KASP markers. The coordinates of SNPs in the TAIR10 and Col-CEN assemblies are provided.

**Table S11. Primer sequences used for *CTL3.9* SSLP and *HS6* dCAPs markers.** The oligonucleotide sequences are provided that were used to; (i) validate T-DNA insertions, or the point mutations in various lines, (ii) to identify SSLPs to map *CEN3* proximal recombination events in wild type, *recq4a recq4b* and *HEI10* overexpression lines, (iii) to fine-map crossovers within *HOTSPOT6* in wild type as derived Cleaved Amplified Polymorphic Sequences (dCAPs) markers, and (iv) primers used to validate T-DNAs in CTL3.9 line.

